# A new lineage of non-photosynthetic green algae with extreme organellar genomes

**DOI:** 10.1101/2021.11.17.468966

**Authors:** Tomáš Pánek, Dovilė Barcytė, Sebastian C. Treitli, Kristína Záhonová, Martin Sokol, Tereza Ševčíková, Eliška Zadrobílková, Karin Jaške, Naoji Yubuki, Ivan Čepička, Marek Eliáš

## Abstract

**Background:** The plastid genomes of the green algal order Chlamydomonadales tend to expand their non-coding regions, but this phenomenon is poorly understood. Here we shed new light on organellar genome evolution in Chlamydomonadales by studying a previously unknown non-photosynthetic lineage. We established cultures of two new *Polytoma*-like flagellates, defined their basic characteristics and phylogenetic position, and obtained complete organellar genome sequences and a transcriptome assembly for one of them.

**Results:** We discovered a novel deeply diverged chlamydomonadalean lineage that has no close photosynthetic relatives and represents an independent case of photosynthesis loss. To accommodate these organisms we establish the new genus *Leontynka*, with two species (*L. pallida* and *L. elongata*) distinguishable through both their morphological and molecular characteristics. Notable features of the colourless plastid of *L. pallida* deduced from the plastid genome (plastome) sequence and transcriptome assembly include the retention of ATP synthase, thylakoid-associated proteins, the carotenoid biosynthesis pathway, and a plastoquinone-based electron transport chain, the latter two modules having an obvious functional link to the eyespot present in *Leontynka*. Most strikingly, the ~362 kbp plastome of *L. pallida* is by far the largest among the non-photosynthetic eukaryotes investigated to date due to an extreme proliferation of sequence repeats. These repeats are also present in coding sequences, with one repeat type found in the exons of 11 out of 34 protein-coding genes, with up to 36 copies per gene, thus affecting the encoded proteins. The mitochondrial genome of *L. pallida* is likewise exceptionally large, with its >104 kbp surpassed only by the mitogenome of *Haematococcus lacustris* among all members of Chlamydomonadales hitherto studied. It is also bloated with repeats, though entirely different from those in the *L. pallida* plastome, which contrasts with the situation in *H. lacustris* where both the organellar genomes have accumulated related repeats. Furthermore, the *L. pallida* mitogenome exhibits an extremely high GC content in both coding and non-coding regions and, strikingly, a high number of predicted G-quadruplexes.

**Conclusions:** With its unprecedented combination of plastid and mitochondrial genome characteristics, *Leontynka* pushes the frontiers of organellar genome diversity and is an interesting model for studying organellar genome evolution.

## Background

Secondary loss of photosynthesis has occurred numerous times across the diversity of plastid-bearing eukaryotes, including land plants [1,2]. Among algae, photosynthesis loss has been most common among groups characterised by secondary or higher-order plastids, with chrysophytes and myzozoans (including apicomplexans as the best-studied non-photosynthetic “algae”) being the most prominent examples. In green algae, loss of photosynthesis is restricted to several lineages within two classes, Trebouxiophyceae and Chlorophyceae [3]. Colourless trebouxiophytes are formally classified in two genera, *Helicosporidium* and the polyphyletic *Prototheca*, collectively representing three independent photosynthesis loss events [4]. While these organisms live as facultative or obligate parasites of metazoans (including humans), non-photosynthetic members of the Chlorophyceae are all free-living osmotrophic flagellates. Two genera of such colourless flagellates have been more extensively studied and are represented in DNA sequence databases: the biflagellate *Polytoma* and the tetraflagellate *Polytomella*. They both fall within the order Chlamydomonadales (Volvocales *sensu lato*), but are not closely related to each other. Furthermore, *Polytoma* as presently circumscribed is polyphyletic, since *P. oviforme* does not group with its congeners, including the type species *P. uvella* [3]. Hence, photosynthesis was lost at least three times in Chlamydomonadales, but the real number is probably higher, as several other genera of colourless flagellates morphologically falling within this group were historically described [5] yet still remain to be studied with modern methods. Indeed, a taxonomically unidentified non-photosynthetic chlamydomonadalean (strain NrCl902) unrelated to any of the three known lineages was reported recently [6]; whether it corresponds to any of the previously formally described taxa has yet to be investigated.

The non-photosynthetic chlamydomonadaleans are not only diverse phylogenetically, but they also exhibit diversity in the features of their residual plastids. Most notably, *Polytomella* represents one of the few known cases of a complete loss of the plastid genome (plastome) in a plastid-bearing eukaryote [7]. In contrast, *Polytoma uvella* harbours the largest plastome amongst all non-photosynthetic eukaryotes studied to date (≥230 kbp). This is not due to it preserving a large number of genes, but because of the massive accumulation of long arrays of short repeats in its intergenic regions [8]. The unusual architecture of the *P. uvella* plastome seems to reflect a more general trend of plastome evolution in Chlamydomonadales, i.e. a tendency to increase in size by the expansion of repetitive sequences. An extreme manifestation of this trend was uncovered by sequencing the 1.35-Mbp plastome of the photosynthetic species *Haematococcus lacustris*, the record-holder amongst all fully sequenced plastomes to date [9,10]. Interestingly, *H. lacustris* also harbours by far the largest known mitochondrial genome (mitogenome) amongst all Chlamydomonadales, which has expanded to 126.4 kbp through the accumulation of repeats highly similar to those found in the plastome, suggesting an inter-organellar transfer of the repeats [11]. The mechanistic underpinnings of the accumulation of repeats in chlamydomonadalean organellar genomes are still not clear.

Whilst studying protists living in hypoxic sediments, we obtained cultures of two colourless flagellates that transpired to represent a novel, deeply separated lineage of Chlamydomonadales. They are herein described formally as two species in a new genus. The sequences of organellar genomes of one of the isolates were determined using a combination of different DNA sequencing technologies, which revealed extreme features concerning their size and/or composition. Our analysis of these genomes provides important new insights into the evolution of organellar genomes in general.

## Results

### A new lineage of non-photosynthetic Chlamydomonadales with two species

Based on their 18S rRNA gene sequences, the two new isolates, AMAZONIE and MBURUCU, constitute a clade (with virtually full bootstrap support) that is nested within Chlamydomonadales, but separate from all the principal chlamydomonadalean clades as demarcated by [12] (Fig. 1a). Notably, this new lineage is clearly unrelated to all previously studied non-photosynthetic chlamydomonadaleans, including *Polytomella* (branching off within the clade *Reinhardtinia*), both lineages representing the polyphyletic genus *Polytoma* (*P. uvella* plus several other species in the clade *Caudivolvoxa*, and *P. oviforme* in the clade *Xenovolvoxa*), and the strain NrCl902 (also in *Caudivolvoxa;* Additional file 1: Fig. S1). The 18S rRNA gene sequences of our strains differed in 13 positions (out of 1,703 available for comparison), and were as deeply diverged in the inferred phylogenetic tree as other chlamydomonadalean pairs classified as separate species or even genera. When compared to the radiations of the other two non-photosynthetic lineages with more than one representative in the tree, the depth of divergence between the two strains was greater than between any pair of species of *Polytoma* (*sensu stricto*), but less than between some of the species pairs of *Polytomella*, a lineage with a highly accelerated rate of 18S rRNA gene evolution (Additional file 1: Fig. S1). In addition, the ITS1-5.8S-ITS2 rDNA regions of the two strains exhibited only 88% identity, with the differences including several compensatory base changes (CBCs) in helix II of the characteristic secondary structure of the ITS2 region (Additional file 1: Fig. S2). This, considered alongside the morphology-based evidence presented below, led to the conclusion that the two strains represent two different species of a new genus of chlamydomonadalean algae, for which we propose the names *Leontynka pallida* (strain AMAZONIE) and *Leontynka elongata* (strain MBURUCU). Formal descriptions of the new taxa are provided in Additional file 2: Note S1 [13–23].

**Fig. 1.**
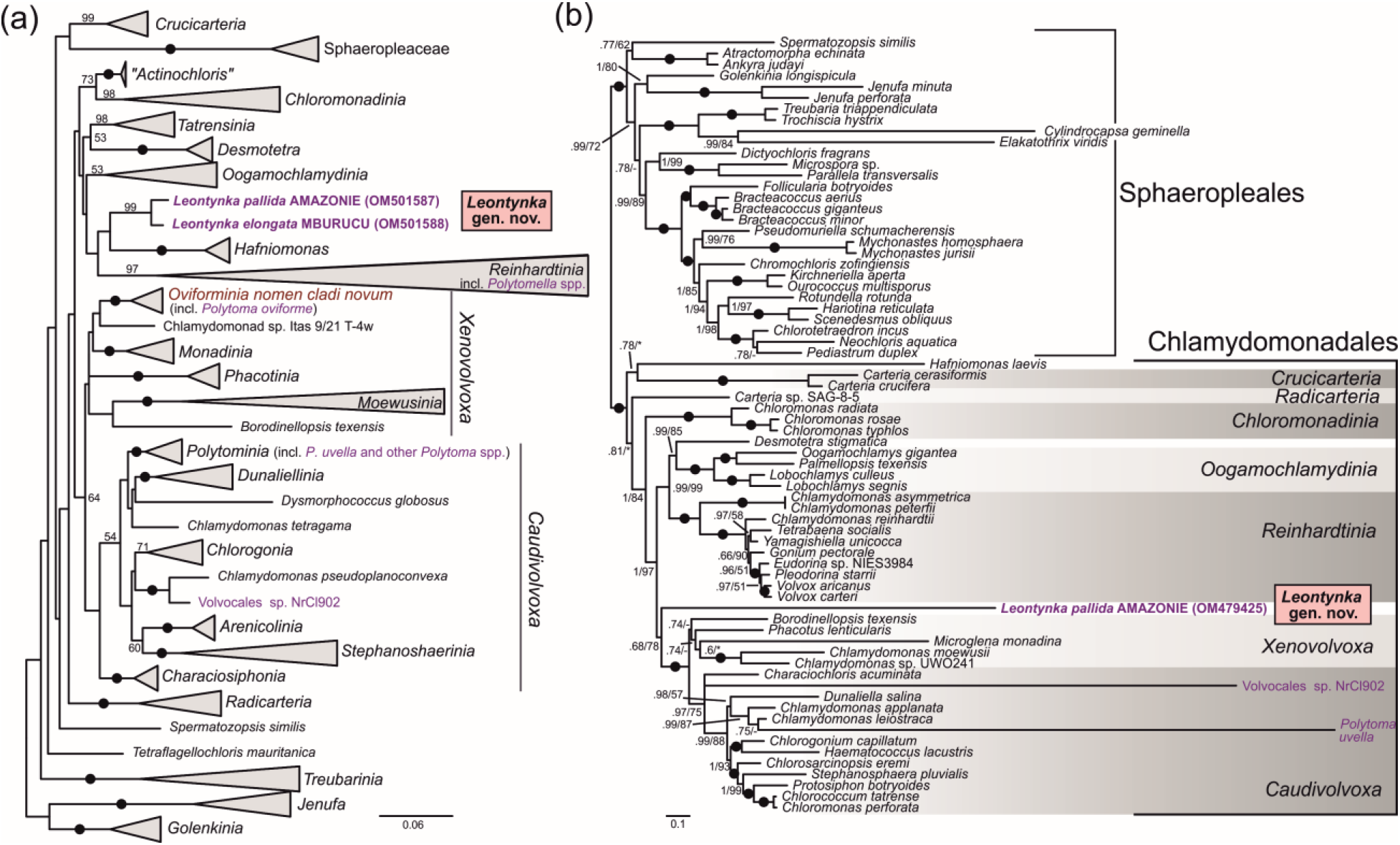
Phylogenetic position of the new genus of non-photosynthetic green algae, *Leontynka*. Non-photosynthetic taxa in Chlamydomonadales are highlighted in violet. (a) Maximum likelihood phylogenetic tree (IQ-TREE, TIM2+F+I+G4 substitution model) of 18S rRNA gene sequences from Chlamydomonadales and related chlorophytes. Non-parametric bootstrap support values calculated from 100 replicates are shown when ≥50. Previously demarcated main clades [12] are collapsed, and outgroup taxa are not shown for simplicity. The full version of the tree is provided as Additional file 1: Fig. S1. (b) Phylogenetic analysis of a concatenated dataset of 24 conserved plastome-encoded proteins (5,020 amino acid positions) from Chlamydomonadales, including *Leontynka pallida*, and the sister order Sphaeropleales (*sensu lato*; see Fučíková *et al*., 2019). The tree topology was inferred using PhyloBayes (CAT+GTR substitution model), branch support values correspond to posterior probability (from PhyloBayes)/maximum likelihood bootstrap analysis (IQ-TREE, LG+C60+F+G4 substitution model, 100 non-parametric bootstrap replicates). Black dots represent full support obtained with both methods, asterisks denote bootstrap support values <50. *Polytomella* spp. and *Polytoma oviforme* are not represented in part b because of the complete absence of a plastid genome and the lack of plastid genome data, respectively.

The phylogenetic position of *L. pallida* was also studied by using protein sequences encoded by its plastome and mitogenome (see below). Phylogenomic analysis of a concatenated dataset of 24 conserved proteins encoded by plastomes of diverse members of Chlorophyceae, including a comprehensive sample of available data from Chlamydomonadales, revealed *L. pallida* to be a separate lineage, potentially sister to a fully supported broader clade comprising representatives of the clades *Caudivolvoxa* and *Xenovolvoxa* (*sensu* [12]; Fig. 1b). This position of *L. pallida* received moderate support from a maximum likelihood analysis (nonparametric bootstrap value of 78%), but inconclusive support from the PhyloBayes analysis (posterior probability of 0.68). Importantly, *L. pallida* was unrelated to *Polytoma uvella* (nested within *Caudivolvoxa* with full support). *Polytoma oviforme* and the genus *Polytomella* were missing from the analysis due to the lack of plastome data and the absence of a plastome, respectively. Consistent with the results of the 18S rRNA gene phylogeny, *Polytomella* and *L. pallida* were supported as unrelated lineages by a phylogenetic analysis of concatenated set of seven mitogenome-encoded proteins (Additional file 1: Fig. S3). This analysis also confirmed that the *Leontynka* lineage represents a lineage falling outside the major chlamydomonadalean clades corresponding to *Reinhardtinia* and *Caudivolvoxa*+*Xenovolvoxa*, but potential relationships to other chlamydomonadalean taxa could not be tested due to the lack of corresponding mitogenome data.

Both *Leontynk*a species lacked a green plastid (chloroplast). Instead, their cells were occupied by a colourless leucoplast containing starch grains, typically filling most of its volume (Fig. 2, Additional file 1: Figs S4 and S5). Two anterior, isokont flagella approximately as long as the cell body emerged from a keel-shaped papilla (Additional file 1: Fig. S4c, d). Cells of both species also contained two apical contractile vacuoles (Fig. 2c, f, Additional file 1: Fig. S4a, c, Fig. S5a, d, h), a central or slightly posterior nucleus (Additional file 1: Fig. S4c, Fig. S5h), inclusions of yellowish lipid droplets (Fig. 2h, Additional file 1: Fig. S5h), and one or occasionally two eyespots (Fig. 2a, b, and f–j, Additional file 1: Fig. S4a, b, e, g, h, Fig. S5a, c–f, h). Reproduction occurred asexually through zoospore formation, typically with up to four zoospores formed per mother cell (Additional file 1: Fig. S4g, h). The two species differed in the cell shape and position of the eyespot, as described in more detail in Additional file 2: Note S2.

**Fig. 2.**
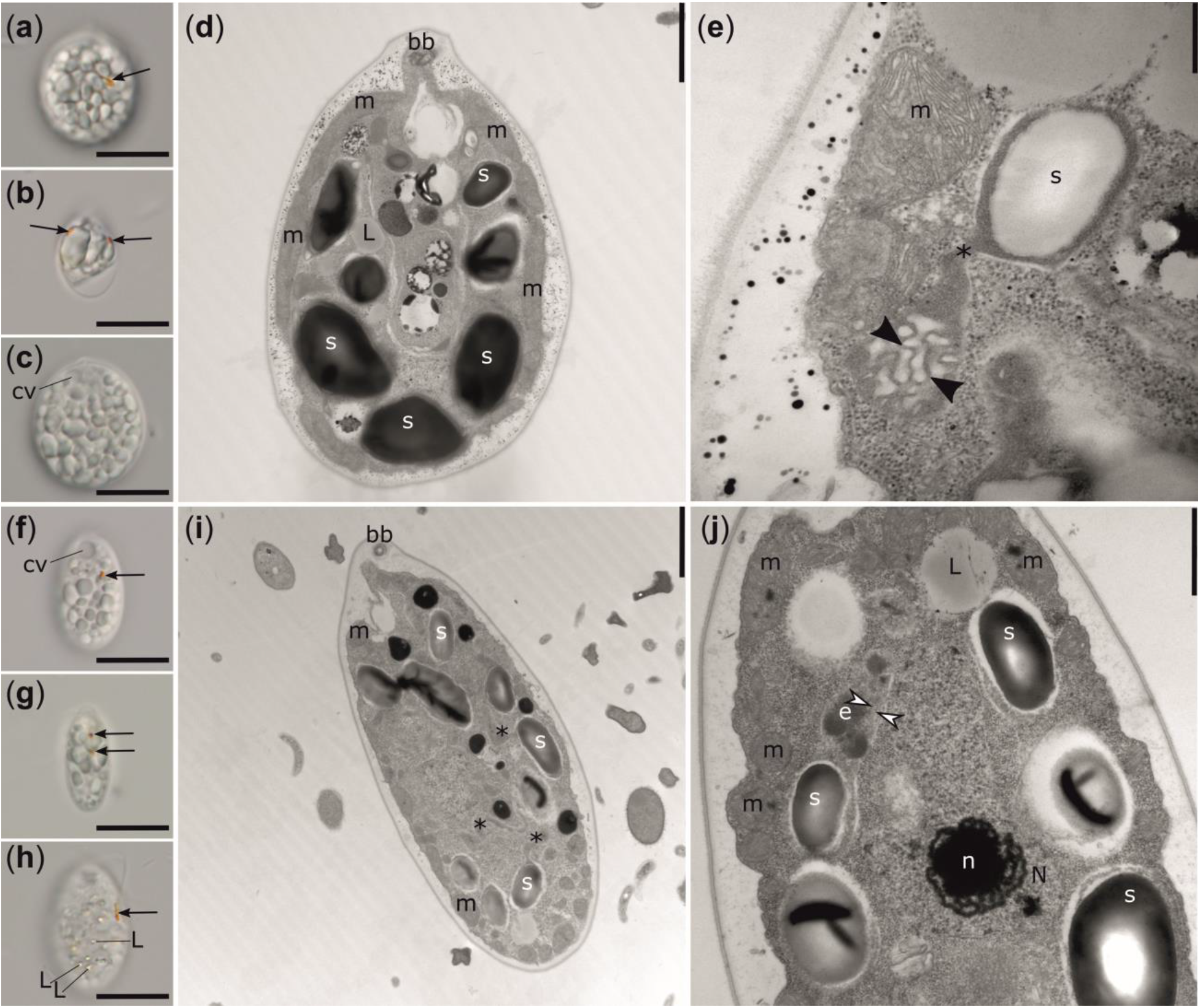
Light and transmission electron microscopy of *Leontynka pallida* (a–e) and *Leontynka elongata* (f–j). Note the difference in the cell shape between the two species and the presence of a single (a, f, h) or two (b, d, g, i, j) eyespots. Lipid droplets were also detected within the cells (d, h, j). Abbreviations: bb – basal body; cv – contractile vacuole; e – eyespot; L – lipid droplet; m – mitochondrion; N – nucleus; n – nucleolus; s – starch. Arrows point to eyespots; asterisks mark plastid “bridges” connecting separate compartments; black arrowheads show membranous inclusions; white arrowheads point to thylakoids. Scale bars: a-c, f-g = 10 μm; d, i = 0.2 μm; e = 0.5 μm; j = 1 μm.

The plastid of both species was bounded by a double membrane and composed of numerous separate compartments connected by narrow “bridges” (Fig. 2d, e, i, j, Additional file 1: Fig. S6a-c, h). Each compartment contained either a single large or two smaller starch grains, leaving essentially no room for the stroma or thylakoids. Rarely, starch-free compartments containing membranous inclusions were present (Fig. 2e). The eyespot globules were inside the plastid and were associated with structures that we interpret as thylakoids (Fig. 2j). Mitochondrial profiles were highly abundant and contained numerous cristae (Additional file 1: Fig. S6e, f, i). It was impossible to unambiguously determine the crista morphology in *L. pallida* (Additional file 1: Fig. S6i), but in *L. elongata*, the cristae were of the discoidal morphotype (Additional file 1: Fig. S6e). Further details on the ultrastructure of *Leontynka* spp. are presented in Additional file 2: Note S2.

### The non-photosynthetic plastid of *Leontynka pallida* has preserved functions commonly associated with photosynthesis

A complete plastome sequence was assembled for *L. pallida* using a combination of Oxford Nanopore and Illumina reads. This was complemented by sequencing and assembling the transcriptome for the same species, yielding 65,552 contigs (summing to a total assembly length of 46,790,231 bp). Despite a filtering step to remove bacterial contamination (see Methods), the transcriptome assembly still contains contaminant sequences, and must therefore be treated with caution. The plastome sequence corresponds to a circular-mapping molecule comprising 362,307 bp (Fig. 3a). Thirty-four protein-coding genes (including two intronic ORFs), 26 tRNA genes (a standard set presumably allowing for translation of all sense codons), and genes for the three standard rRNAs were identified and annotated in the genome. Three genes are interrupted by introns (with the splicing verified by transcriptome data): *atpA* with one group I intron that contains an ORF encoding a LAGLIDADG homing endonuclease, *tufA* with one group II intron that contains an ORF encoding a reverse transcriptase/maturase protein, and *rnl* with one group I and one group II intron, neither containing an ORF. No putative pseudogenes or apparent gene remnants were identified in the *L. pallida* plastome.

**Fig. 3.**
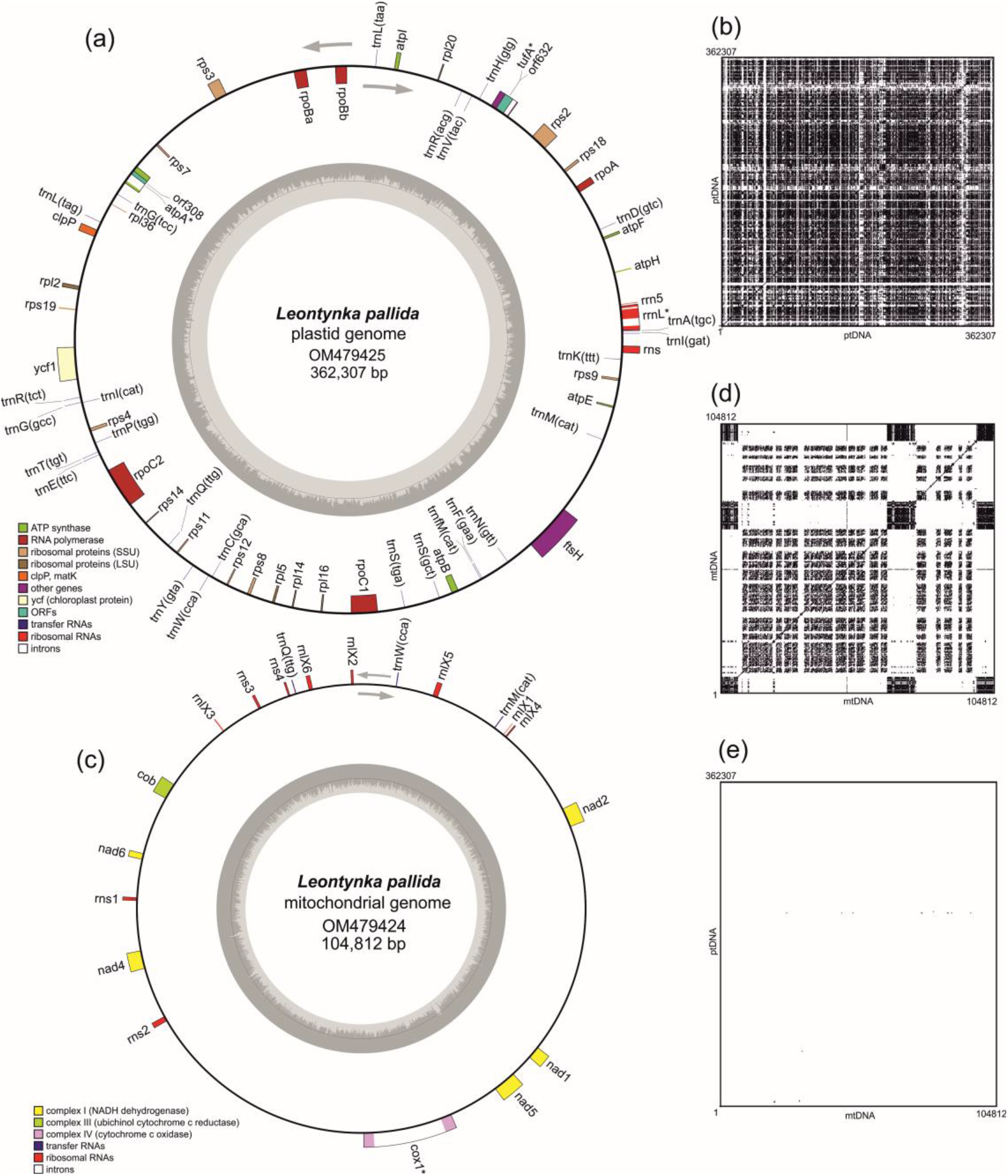
Characteristics of the organellar genomes of *Leontynka pallida*. (a) Gene map of the *L. pallida* plastid genome. Genes are shown as squares (coloured according to the functional category; see the graphical legend in the bottom left corner) on the inner or outer side of the outer circle depending on their orientation (transcription in the clockwise or counter-clockwise direction, respectively; see the grey arrows). Genes marked with an asterisk contain introns. The inner circle represents a GC content plot. (b) Sequence self-similarity plot of the *L. pallida* plastome (ptDNA). (c) Map of the *L. pallida* mitochondrial genome. The display convention is the same as for the plastid genome. (d) Sequence self-similarity plot of the *L. pallida* mitochondrial genome (mtDNA). (e) *L. pallida* plastome-mitogenome similarity plot. All similarity plots were generated by using a word size of 15 bp and black dots represent the occurrence of the same word at the places compared.

No genes that encode proteins directly associated with photosynthetic electron transport components or CO2 fixation were identified in the *L. pallida* plastome. The retained genes encode proteins involved in transcription (RNA polymerase subunits), translation (*tufA* and ribosomal subunit genes), protein turnover (*clpP, ftsH*), and a protein without a clearly defined function (*ycf1*). Nearly all these genes have also been preserved in the plastome of *P. uvella* [8], except for *rps2*. The two non-photosynthetic chlamydomonadalean species share the absence of genes for two ribosomal proteins: *rpl32*, which is also missing from a subset of photosynthetic representatives of Chlamydomonadales, and *rpl23*, which has been conserved in the plastomes of all photosynthetic chlorophytes investigated to date [24]. Though an Rpl32 protein with a predicted plastid-targeting presequence is encoded by the *L. pallida* nuclear genome (Additional file 3: Table S1), the loss of *rpl23* does not seem to be compensated in a similar way. Interestingly, *rpl23* was also independently lost from the plastome of *Helicosporidium* sp. [8], which suggests that this ribosomal subunit may become dispensable upon the loss of photosynthesis. Unlike *rpl32* and similar to *rpl23*, none of the other genes present in the plastome of the reference species *Chlamydomonas reinhardtii* but absent from the *L. pallida* plastome has been complemented by a nuclear version, as indicated by our analyses of the *L. pallida* transcriptome assembly.

The most notable feature of the *L. pallida* plastid gene repertoire is the preservation of genes encoding ATP synthase subunits, i.e., *atpA, atpB, atpE, atpF, atpH*, and *atpI*. It is the same set of ATP synthase subunit genes as found in the plastomes of photosynthetic green algae; conversely, the plastome of *P. uvella* has none of these genes. As evidenced by the transcriptome data, the three missing ATP synthase subunits (AtpC, AtpD, and AtpG) are encoded by the *L. pallida* nuclear genome and bear predicted plastid-targeting signals (Additional file 3: Table S1). It was proposed that the retention of ATP synthase in certain non-photosynthetic plastids is functionally linked to the retention of the twin-arginine protein translocase (Tat; [25]), and indeed we found homologs of all three Tat subunits (TatA, TatB, TatC) in the nuclear transcriptome of *L. pallida* (Additional file 3: Table S1). The ATP synthase and Tat translocase are both normally localised to thylakoid membranes, and we could discern putative thylakoids associated with the eyespot in *Leontynka* (see above). Consistent with these considerations, we identified the *L. pallida* transcriptome assembly homologs of components of several additional thylakoid-associated protein-targeting or translocation systems [26,27,28], namely the plastidial SRP pathway (cpSRP54 and cpFtsY), ALB3 protein insertase, and thylakoid-specific Sec translocase (Additional file 3: Table S1). Furthermore, in *L. pallida* we also found homologs of proteins implicated in thylakoid biogenesis, such as VIPP1, FZL, THF1, and SCO2 ([29]; Additional file 3: Table S1).

The reddish colour of the *Leontynka* eyespots suggests the presence of carotenoids (similar to the eyespot of *C. reinhardtii* [30]). In addition, searches of the *L. pallida* transcriptome assembly revealed the presence of a homolog of the *C. reinhardtii* eyespot-associated photosensor channelrhodopsin 1 (ChR1; Additional file 3: Table S1) that requires a carotenoid derivative, retinal, as a chromophore [31]. Indeed, our analysis of the *L. pallida* transcriptome assembly revealed the presence of enzymes of a plastid-localised carotenoid biosynthetic pathway that is common in photosynthetic species (Additional file 3: Table S1). Notably, *L. pallida* has also retained enzymes for the synthesis of plastoquinone, which serves as an electron acceptor in two reactions of carotenoid biosynthesis, and the plastid terminal oxidase (PTOX), which recycles plastoquinone (from its reduced form plastoquinole) by further passing the electrons to molecular oxygen ([6]; Additional file 3: Table S1). Altogether, genomic and transcriptomic evidence confirms the loss of photosynthesis in *L. pallida* deduced through microscopic observation, yet concurrently indicates the preservation of various plastidial functions that are commonly viewed as primarily serving the photosynthetic role of the organelle.

### The *L. pallida* plastome is bloated due to extreme expansion of sequence repeats

Above all the aforementioned interesting features of the plastid gene repertoire, what makes the *L. pallida* plastome truly peculiar are the intergenic regions. Their average length is 4.7 kbp, which is 1.5 and five times greater than the average length of the intergenic regions in the plastomes of *P. uvella* and *C. reinhardtii*, respectively (Table 1). Furthermore, while the GC content of the *P. uvella* intergenic regions is vastly different from that of coding regions (19% versus 40%), the GC content of these two plastome partitions is highly similar in *L. pallida* (as well as in *C. reinhardtii*; Table 1). A self-similarity plot generated for the *L. pallida* plastome revealed massive repetitiveness of the DNA sequence, with only short islands of unique sequence scattered in the sea of repeats (Fig. 3b). The repeats are highly organised and occur in various arrangements: tandem repeats, interspersed repeats, inverted repeats (palindromes), and other higher-order composite repeated units. As an example, let us take the most abundant repeat, the imperfect palindrome (IP) CAAACCAGT|NN|ACTGGTTAG. More than 1,300 copies are present, with the dinucleotide AA as the predominant form of the internal spacer. This repeat is mostly localised in clusters (>1,200 cases) where its copies are interleaved with a repeat with the conserved sequence TAACTAAACTTC, so together they constitute an extremely abundant composite tandem repeat. In a single region, the palindromic repeat combines with a different interspersed repeat (TAACTACTT), together forming a small cluster of composite tandem repeats with 14 copies. Additionally, the same palindromic repeat is part of another, 146 bp-long repeat present with 27 copies across the plastome (for details see Additional file 2: Note S3).

**Table 1.** Basic characteristics of plastomes of *Leontynka pallida*, selected other non-photosynthetic chlorophytes, and the photosynthetic *Chlamydomonas reinhardtii* and *Haematococcus lacustris* for comparison.

| species | plastomes |  | genes |  | coding DNA** |  | intergenic regions |  |  |
| --- | --- | --- | --- | --- | --- | --- | --- | --- | --- |
|  | accession number | size (bp) | protein-coding* | tRNAs | GC (%) | total length (%) | GC (%) | average length (bp) | total length (%) |
| *Leontynka pallida* | OM479425.1 | 362,307 | 32 | 26 | 35 | 18 | 37 | 4,726 | 80 |
| *Polytoma uvella* | KX828177.1 | 230,207 | 25 | 27 | 40 | 20 | 19 | 2,998 | 68 |
| *Volvocales* sp. NrCl902 | LC516060.1 | 176,432 | 25 | 30 | 36.1 | 32.6 | 42.8 | 1,944 | 60.9 |
| *Helicosporidium* sp. | DQ398104.1 | 37,454 | 26 | 25 | 27.6 | 95.4 | 13.4 | 33.8 | 4.4 |
| *Chlamydomonas reinhardtii* | NC_005353.1 | 203,828 | 69 | 29 | 34.6 | 43 | 34 | 923 | 48.5 |
| *Haematococcus lacustris* (†) | NC_037007.1 | 1,352,306 | 63 | 28 | 38.6 | 6.8 | 51 | 12,326 | 91.15 |

\*does not include ORFs inside introns
\*\*does not include introns and ORFs inside introns, but includes tRNAs and rRNAs
(†) since the annotation of the *H. lacustris* plastome available in the respective GenBank record is highly incomplete, the values presented are based on a reannotation obtained using MFannot (with a genetic code translating UGA as Trp, based on the insights of [10]).

Apart from intergenic regions, sequence repeats are also found in the four introns present in the *L*. *pallida* plastome. The most prominent is a cluster of 20 copies of the motif TGGTTAGTAACTAAACTTCCAAACCAGTAAAC in the intron inside the *atpA* gene that is also abundant in intergenic regions (more than 1,000 copies typically located in huge clusters). Strikingly, when analysing the distribution of the most abundant IPs, we noticed that the motif AAGCCAGC|NNN|GCTGACTT and its variants are also present in coding regions, namely in the exons of 11 out of 34 protein-coding genes of the *L*. *pallida* plastome (Fig. 4a-c). They can be present in numbers up to 36 copies per gene (“variant 8” in exons of *rpoC2;* Fig. 4b). In most cases the IP motif is part of a longer repeat unit that includes extra nucleotides at both ends (“variant 4” to “variant 8” in Fig. 4c). The most complex repeat unit variant is the following one (the IP core in round brackets; square brackets indicate alternative nucleotides occurring at the same position): AAAGAT-(AAGTCAGC|AGA|GCTGAC[AT]T)-CCAGACCACTAAAGTGGTCAGTAACTAAAAGTTAT. It is restricted to coding sequences (i.e., is absent from intergenic regions and introns) and occurs in eight copies inside three genes (*rpoC2, rpoC1, ftsH*), resulting in an insertion of a stretch of 20 amino acid residues in the encoded proteins. Other repeat variants (listed in Fig. 4c) have proliferated in exons as well as intergenic regions and introns. However, not one of the aforementioned nucleotide motifs was found in plastomes of other chlamydomonadalean algae, indicating they have originated and diverged only in the *Leontynka* lineage.

**Fig. 4.**
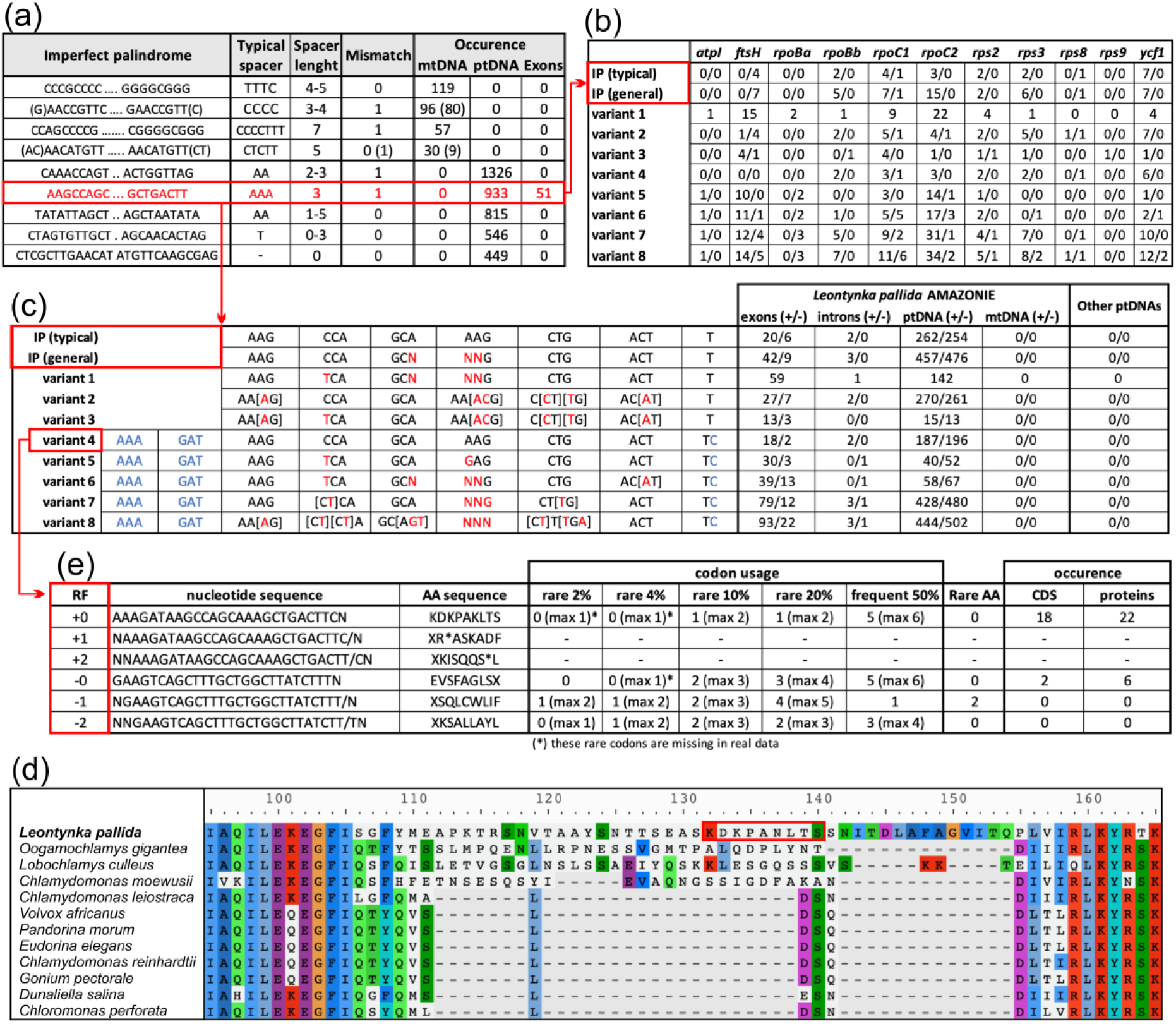
Distribution of repeats in organellar genomes of *Leontynka pallida*. (a) The most abundant imperfect palindromes and their characteristics. The “Spacer” corresponds to the presumed loop separating the palindromic regions that presumably pair to form a stem structure. The “Mismatch” column indicates the number of positions that deviate from a perfect palindrome. The occurrence of the repeats is given for the plastome (ptDNA) and mitogenome (mtDNA), with the number of cases indicated for the whole organellar genome, and separately for exons in protein-coding genes. In two cases of mitogenome repeats, two variants – a shorter and a longer – are considered, with the latter indicated in parentheses. (b) Distribution of the imperfect palindrome AAGCCAGC|NNN|GCTGACTT and its most common variants within exons of the plastome. The numbers show the abundance of the given repeat in the direct/reverse complement orientation (relative to the coding sequence). In the case of “variant 1”, the repeat has the same sequence in both directions, so only one number per gene is presented. Note that the variants considered are not mutually exclusive alternatives, but correspond to nested categories with a different degree or relaxation of the sequence pattern. (c) Characterisation of repeats from (b) and their abundance in various regions of the plastome and the mitogenome of *L*. *pallida* as well as other plastomes of Chlamydomonadales deposited in NCBI databases. The numbers show the abundance of the given repeat in the direct/reverse complement orientation (relative to the coding sequence in the case of exons, or relative to the DNA strand corresponding to the reference organellar sequence in the case of the values for the whole organellar genome). (d) Occurrence of the “variant 8” repeat (translated in the reading frame +0 as KDKPANLTS) in a variable region of the ribosomal protein Rps8 (detail; the full alignment is available as Additional file 1: Fig. S9). (e) Occurrence of the “variant 4” repeat in protein-coding sequences and its translation for all six reading frames. The category of rare codons (“rare 2%”) is defined as the sum of the least used codons, together representing less than 2% of all codons in the plastome (100% = 19,899 codons); the categories of the 4%, 10%, and 20% rarest codons and that of more than 50% of the most frequent codons are defined following the same convention (listed in Additional file 3: Table S5). The numbers indicated for “codon usage” correspond to the minimum number of the codons of the respective category present in the respective reading frame, with the “max X” numbers indicating the maximum number of such codons, depending on the actual nucleotide sequence of the degenerated “variant 4” repeat. Note that some rare codons (2-4% category) are not observed in the actual *L. pallida* plastid gene sequences, although their presence would be theoretically possible (see the asterisks). The column “Rare AA” indicates the occurrence of amino acids belonging to the category of amino acids generally rarely used in plastome-encoded proteins in *L. pallida* (see Additional file 3: Table S6). The occurrence of the repeat variants indicated for coding sequences (CDS) corresponds to their occurrence as counted at the nucleotide level, whereas the occurrence in proteins is counted at the amino acid sequence level (and may be higher due to different nucleotide sequences encoding the same amino acid sequence). The analysis of intraexonic repeat insertions is discussed in more detail in Additional file 2: Note S4.

Manual inspection of protein sequence alignments including chlamydomonadalean orthologs of the *L. pallida* proteins revealed that the intraexonic repeat insertions are located mostly in poorly conserved regions (see Fig. 4d for an example). Inspection of RNA-seq reads mapped onto the plastome sequence did not reveal any region with significantly lower read coverage (which would indicate the presence of hidden introns in predicted repeats-containing exons). Preferential proliferation in variable parts of coding regions is consistent with a high abundance of these motifs in proteins that exhibit a general tendency for including rapidly evolving and poorly conserved regions, namely FtsH (Additional file 1: Fig. S7), Ycf1, RpoC1, RpoC2, and RpoBb. Interestingly, the phase and orientation of the intraexonic repeats is not random with respect to the reading frame and the direction of transcription, and is potentially biased such that not only termination codons, but also codons generally rare in *L. pallida* plastid coding sequences are avoided from the actual frame in which the insertion is read during translation (for details see Fig. 4c, e, Additional file 2: Note S4). This bias does not merely reflect a possible bias in the orientation of the repeats relative to the DNA strand of the genome, as the repeats are distributed roughly equally in both strands when counted at the whole-genome level (Fig. 4c).

### The GC-rich mitogenome of *Leontynka pallida* contains a high number of potential quadruplex-forming sequences

The mitogenome sequence was assembled from Nanopore and Illumina reads as a linear molecule of 110,515 bp with long (~5,770 bp) nearly perfect (97.7% identity) direct terminal repeats differentiated primarily by the presence/absence of two short repetitive regions (13 and 70 bp) (Fig. 3c, Additional file 1: Fig. S8). This suggests that the *L. pallida* mitogenome is in fact circular, with the slight differences in the terminal direct repeats of the assembled linear contigs reflecting sequence variability of a particular genomic region between the different genome copies in *L. pallida*, or possibly sequencing or assembly artefacts. If circular, the mitogenome would then have a length of ~104,812 bp. The suspected circularity of the mitogenome is also compatible with the absence of the *rtl* gene, which is present in all linear mitogenomes of Chlamydomonadales characterised to date and encodes a reverse transcriptase-like protein implicated in the replication of the mitogenome termini [32]. Apart from *rtl*, the gene content of the *L. pallida* mitogenome is essentially the same as in other previously sequenced chlamydomonadalean mitogenomes, and includes seven protein-coding genes (with *cox1* interrupted by an ORF-free group II intron), only three tRNA genes, and regions corresponding to the 16S and 23S rRNA genes. As in other chlamydomonadaleans studied in this regard [33–35], the mitochondrial 16S and 23S rRNA genes in *L. pallida* are fragmented, consisting of multiple separately transcribed pieces. Four fragments, together constituting a presumably complete 16S rRNA, were annotated by considering the sequence and secondary structure conservation of the molecule. The number of the 16S rRNA fragments is thus the same as in *Chlamydomonas reinhardtii*, but the breakpoints are not. Due to lower conservation of the 23S rRNA molecule, we could identify only a few of the presumed gene fragments in the *L. pallida* mitogenome.

The large size and the low density of coding sequences in the *L. pallida* mitogenome (~84.8% of its complete sequence is represented by intergenic regions) are atypical for Chlamydomonadales, including the other non-photosynthetic species: the mitogenome of *P. uvella* is 17.4 kbp long [36], and in *Polytomella* spp. the mitogenome size ranges from ~13 to 24.4 kbp [37–38]. In fact, the *L. pallida* mitogenome is comparable only to the recently characterised mitogenome of *Haematococcus lacustris*, which, with the same gene content, is even larger (126.4 kbp) yet with a similar representation of intergenic regions (83.2%). A self-similarity plot generated for the *L. pallida* mitogenome revealed a highly repetitive nature of its genome sequence (Fig. 3d), similar to the plastome. However, the repeats are distributed less evenly than in the plastome, being present particularly in the terminal regions of the assembled linear sequence and in several internal hotspots.

With a GC content of 62.6% (as counted for the circularised version of the genome), the mitogenome of *L. pallida* has the third highest documented mitochondrial GC content out of the 11,077 examined mitogenomes available in GenBank, being surpassed only by the lycophyte *Selaginella moellendorffii* (68.2%; [39–40]) and the green alga *Picocystis salinarum* (67.7%). These values contrast sharply with the median GC content value (38%) for the whole set of the mitogenomes examined. We also encountered an exceptionally high GC content (63.3%) and a strong bias towards using GC-rich codons in all protein-coding genes in the *L. pallida* mitogenome (see Additional file 3: Tables S2 and S3). Only two organisms are presently known to have an even higher GC content of their mitochondrial protein-coding genes: the sponge *Leucosolenia complicata* (71.2%; [41]) and *Picocystis salinarum* (67.9%). Some *L. pallida* mitogenome-encoded proteins, namely Nad2 and Nad5, also exhibit a higher relative content of amino acids with GC-rich codons (G, A, R, and P) compared to most of their orthologs in other species (Additional file 3: Table S4). Thus, not only the expanded GC-rich intergenic regions, but also coding regions of the *L*. *pallida* mitogenome contribute to its extremely high GC content.

The repeats in the plastome and mitogenome of *H. lacustris* are nearly identical [11], so it was interesting to compare the two *L. pallida* organellar genomes to find out whether they behave similarly. However, as follows from the respective similarity plot (Fig. 3e) and comparison of the most abundant inverted repeats and palindromes (Fig. 4a), the repeats in the two genomes do not resemble each other. The proliferation of different repeats in the two organellar genomes of *L*. *pallida* at least partially accounts for their strikingly different GC content (62.6% versus 37%). Interestingly, the most abundant IP in the *L*. *pallida* mitogenome contains the GGGG motif (Fig. 4a), which prompted us to bioinformatically investigate the possible occurrence of G-quadruplexes, i.e. unusual secondary structures in nucleic acids formed by guanine-rich regions [42]. Indeed, the *L. pallida* mitogenome was suggested to include up to 14.7 potential quadruplex-forming sequences (PQS) per 1,000 bp. A similar value was inferred for the *S. moellendorffii* mitogenome (15.6 PQS per 1,000 bp), whereas the other mitochondrial and plastid genomes that we analysed for comparison (for technical reasons focusing on GC-rich genomes only) exhibited a much lower values (0.0-6.9 PQS per 1,000 bp; see Additional file 3: Table S2).

## Discussion

The 18S rRNA, plastid, and mitochondrial gene sequence data all support the conclusion that the two strains investigated in this study, AMAZONIE and MBURUCU, represent a phylogenetically novel lineage within Chlamydomonadales that is unrelated to any of the previously known non-photosynthetic lineages in this order, i.e. *Polytomella, Polytoma sensu stricto* (including the type species *P. uvella*), *Polytoma oviforme*, and the recently reported strain NrCl902. However, morphological features of AMAZONIE and MBURUCU, including the cell shape and the presence of two flagella, papilla, an eyespot, and starch granules, make our organisms highly reminiscent of the genus *Polytoma* [5]. This is consistent with the previous insight that the *Polytoma* morphotype does not define a coherent phylogenetic unit [3]. All other historically described genera of colourless flagellates assigned formerly to Chlamydomonadales are sufficiently different from our strains to consider them a potential taxonomic home for AMAZONIE and MBURUCU (see Additional file 2: Note S5), justifying the erection of the new genus *Leontynka* to accommodate the two strains. Furthermore, these strains clearly differ from each other in morphology (including cell shape and size, and position of the eyespot), and are genetically differentiated, as is apparent from the comparison of the 18S rRNA gene and ITS2 region sequences. Indeed, given the presence of several CBCs in helix II of the conserved ITS2 secondary structure, the two strains are predicted to be sexually incompatible, and hence representing separate “biological species” [43–44]. We considered the possibility that AMAZONIE and MBURUCU may represent some of the previously described *Polytoma* species, but as detailed in Additional file 2: Note S5, none seems to be close enough in morphology, as reported in the original descriptions. Given the fact that the majority of *Polytoma* species have been isolated and described from central Europe, whereas our strains both come from tropical regions of South America, it is not so surprising that we have encountered organisms new to science.

*Leontynka* spp. exhibit a number of ultrastructural similarities to the previously studied *Polytoma* species [45–47]. Previous ultrastructural studies of *Polytoma obtusum* [46] and *Polytomella* sp. [48] showed that their mitochondria possess lamellar and irregular tubulo-vesicular cristae, respectively. The cristae of *L. pallida* resemble the latter morphotype, whereas *L. elongata* most probably possesses discoidal cristae (Additional file 1: Fig. S6e, f). Discoidal cristae are a very rare morphotype within the supergroup Archaeplastida, although they apparently evolved several times independently during eukaryote evolution [49] and were previously noticed in several other non-photosynthetic chlorophytes (*Polytoma uvella, Polytomella agilis*, and *Prototheca zopfii*; [50]).

A particularly notable feature of *Leontynka* spp. is the presence of two eyespots. These were more frequent in *L. elongata* (about half of the examined cells had two eyespots), whereas in the *L. pallida* cultures, such cells were rather rare. Variation in the number of eyespots (from none to multiple) in *Chlamydomonas reinhardtii* was shown to be a result of genetic mutations [51], but the factors behind the eyespot number variation observed in *Leontynka* spp. are unknown. An additional unanswered open question is that of what the actual physiological function of the eyespot in a non-photosynthetic organism is. It seems that light frequently remains an important environmental cue for chlamydomonadaleans, even in the absence of photosynthesis, as evidenced by the preservation of an eyespot that is also observable in some species of the genera *Polytoma* and *Polytomella* and in the strain NrCl902 [5,6]. Notably, like *L. pallida*, and with an obvious functional connection to the eyespot, the other non-photosynthetic chlamydomonadaleas that were investigated in this regard have also preserved an intact plastid-localized carotenoid biosynthetic pathway [6,52]. The retention of carotenoid biosynthesis, specifically the enzymes 15-cis-phytoene desaturate and zeta-carotene desaturase, is the obvious factor explaining the preservation of plastoquinone biosynthesis and plastid terminal oxidase (PTOX), which *L. pallida* shares with the *Polytomella* species and the strain NrCl902 [6].

The eyespot in *Leontynka* (shown for *L. elongata* in Fig. 2j) is associated with the structure that we have interpreted as a thylakoid. Indeed, in the well-studied cases of *C. reinhardtii* and some other chlamydomonadalean algae, the layers of pigment granules are organized on the surface of thylakoids closely apposed to the plastid envelope [30, 53]. The existence of thylakoids in *L. pallida* is supported by our identification in the transcriptome assembly of a comprehensive set of proteins underpinning the formation of, or physically associated with thylakoids (Additional file 3: Table S1). Interestingly, some of the corresponding transcripts have very low read coverage, or are even represented by incomplete sequences, suggesting a low level of gene expression and presumably low abundance of the respective proteins. These observations support the notion that the thylakoid system is preserved in *Leontynka* plastids, though in an inconspicuous and likely reduced form. Nevertheless, we cannot rule out that at least some of these proteins or complexes may have relocalised to the inner bounding membrane of the *Leontynka* plastid, or even to a cellular compartment other than the plastid (as suggested for some of these proteins by the results of *in silico* targeting prediction; Additional file 3: Table S1). The exact localisation of these complexes, the actual substrates of the plastidial SRP pathway, ALB3 insertase, and Tat and Sec translocases, and indeed, the physiological functions of the *L. pallida* leucoplast as a whole remain subjects for future research. Despite these uncertainties, the data presented for *Leontynka* clearly make it an independent case supporting the notion that retention of the eyespot constrains the reductive evolution of non-photosynthetic plastids.

*Leontynka* is significant not only as a novel non-photosynthetic group *per se*, but also as an independent lineage within Chlamydomonadales lacking any close photosynthetic relatives. Specifically, based on the phylogenetic analysis of plastome-encoded proteins, *Leontynka* branches off between two large assemblages, each comprised of several major chlamydomonadalean clades defined by [12]. One of these assemblages (potentially sister to *Leontynka*) is comprised of the *Caudivolvoxa* and *Xenovolvoxa* clades, and the other includes *Reinhardtinia, Oogamochlamydinia*, and the genus *Desmotetra* (Fig. 1a). The radiation of the *Reinhardtinia* clade itself was dated to ~300 MYA [54], so the last common ancestor of *Leontynka* and any of its presently known closest photosynthetic relatives must have existed even earlier. In other words, it is possible that *Leontynka* has been living without photosynthesis for hundreds of millions of years. The loss of photosynthesis in the four other known colourless chlamydomonadalean lineages was certainly not that far in the past. Specifically, the origin of *Polytomella* must postdate the radiation of *Reinhardtinia*, owing to the position of the genus with this clade, whereas *Polytoma sensu stricto* (*P. uvella* and relatives) has close photosynthetic relatives (*Chlamydomonas leiostraca, C. applanata* etc.) within the clade *Polytominia* in *Caudivolvoxa* (Fig. 1, Additional file 1: Fig. S1). *Polytoma oviforme* is specifically related to the photosynthetic *Chlamydomonas chlamydogama*, together constituting a clade in *Xenovolvoxa* that has not been formally recognised before, and which we here designate *“Oviforminia”* (Additional file 1: Fig. S1). Finally, the recently reported strain NrCl902 is closely related to the photosynthetic *Chlamydomonas pseudoplanoconvexa* (Fig. 1A; Additional file 1: Fig. S1). The independent phylogenetic position of *L. pallida* based on plastome-encoded proteins is unlikely to be an artefact stemming from the increased substitution rate of *L. pallida* plastid genes that is manifest from the markedly longer branch of *L. pallida* in the tree compared to most other species included in the analysis. Indeed, the branches of *P. uvella* and the strain NrCl902 are even longer (Fig. 1B), yet both organisms are placed at positions consistent with the 18S rRNA gene tree (Fig. 1A; Additional file 1: Fig. S1). Nevertheless, whether *Leontynka* represents a truly ancient non-photosynthetic lineage, or whether it diverged from a photosynthetic ancestor rather recently needs to be tested by further sampling of the chlamydomonadalean diversity, as we cannot rule out the possibility that photosynthetic organisms closely related to the genus *Leontynka* will be discovered.

The considerations presented above concerning the different ages of the separately evolved non-photosynthetic chlamydomonadalean lineages are somewhat at odds with features of their plastomes. Despite the presumably more recent loss of photosynthesis as compared to *Leontynka*, both *P. uvella* and the strain NrCl902 exhibit a more reduced set of plastid genes (Table 1), whereas in *Polytomella*, plastome reduction triggered by photosynthesis loss has reached its possible maximum, i.e., a complete disappearance of the genome. It is likely that factors other than evolutionary time are contributing to the different degrees of plastome reduction in different evolutionary lineages, although little is known in this regard. Compared to *P. uvella* and the strain NrCl902, *L. pallida* has preserved one gene for a plastidial ribosomal protein (*rps2*), and intriguingly, all standard plastidial genes for ATP synthase subunits, complemented by three further subunits encoded by the nuclear genome to allow for the assembly of a complete and presumably functional complex. It was previously hypothesised that ATP synthase is retained by certain non-photosynthetic plastids to generate (at the expense of ATP) a transmembrane proton gradient required for the functioning of Tat translocase [25]. Our identification of all three Tat subunits in *L. pallida* thus provides further support to this hypothesis. However, it must be noted that certain members of the non-photosynthetic trebouxiophyte genus *Prototheca* possess plastidial ATP synthase in the absence of Tat translocase [4], suggesting that ATP synthase may be retained by a non-photosynthetic plastid for other roles than supporting the function of the Tat system. One such possibility, discussed by [55], is mediating the control of the stromal pH (by pumping protons out) to maintain a basic level optimal for the function of certain stromal enzymes. This hypothesis should be tested in the future by comparing repertoires of plastidial enzymes reconstructed for multiple non-photosynthetic representatives possessing and lacking the plastidial ATP synthase. The role of ATP synthase in *L. pallida* in the functioning of the eyespot as hypothesized for *C. reinhardtii* [56] could also be directly relevant for its retention.

The most intriguing feature of *L. pallida* is the extreme expansion of its organellar genomes. Generally, organellar genomes exhibit remarkable variation in gene content, architecture, and nucleotide composition, with most of them being AT-rich. With its GC content of only ~37%, the *L. pallida* plastome is no exception in this respect. As noted by [10], 98% of plastomes are under 200 kbp and harbour modest amounts (<50%) of non-coding DNA. The *L. pallida* plastome, reaching 362.3 kbp, may not seem that impressive in comparison with the giant plastomes recently reported from some photosynthetic species, including the distantly related chlamydomonadalean *Haematococcus lacustris* (1.35 Mbp; [9]) or certain red algae (up to 1.13 Mbp; [57]); however, it by far dwarfs the plastomes of all non-photosynthetic eukaryotes studied to date. The previous record holder, the ~230 kbp plastome of *Polytoma uvella* [8], is only two thirds of the size of the *L. pallida* plastome. The difference is not only because of the higher number of genes, but primarily because of a more extreme expansion of intergenic regions in *L. pallida* (4.7 kbp on average) than in *P. uvella* (3.0 kbp on average; Table 1). The plastome of the strain NrCl902 with its size of 176.4 kbp, despite exhibiting the same gene content as the *P. uvella* plastome, is much less extreme [6], although still with the intergenic regions substantially expanded as compared to the plastomes of non-photosynthetic trebouxiophytes (Table 1).

Thus, despite its uniqueness, the organisation of the *L. pallida* plastome fits into the general pattern observed in chlamydomonadalean algae, where plastomes in different lineages tend to increase in size by accumulating repetitive sequences [10,58]). It has been suggested that the repeats are prone to double-strand breaks, which are then repaired by an error-prone mechanism favouring repeat expansion [59]. However, the plastome of *L. pallida* is bloated not only due to the extreme proliferation of repetitive DNA in intergenic regions, but also due to the expansion of some of them into the intronic regions and, much more surprisingly, even into exons (Fig. 4). The biased orientation and phase of the insertions with respect to the coding sequence and the reading frame avoid introduction of termination codons as well as rare codons or codons for rare amino acids (C, W) into the coding sequences (Fig. 4 c, d, Additional file 2: Note S4), which suggests that purifying selection eliminates those insertions that would disrupt or reduce the efficiency of translation of the respective mRNAs. Still, exons provide an important niche for the repeats: for example, for the “variant 8” repeat, the exonic copies constitute ~12.2% of the whole repeat population (compared to protein-coding sequences constituting ~17.2% of the total plastome length). Such a massive proliferation of repeats to coding regions is unprecedented to our knowledge, although a much less extensive invasion of a different repeat into coding sequences was recently noticed in the plastome of another chlamydomonadalean alga, *Chlorosarcinopsis eremi* [59]. Here the repeats are found in small numbers in the genes *ftsH, rpoC2*, and *ycf1*, paralleling the situation in *L. pallida* and consistent with the notion that genes encoding proteins that are rich in poorly conserved regions are more likely to tolerate the invasion of the repeats.

Recent sequencing of the mitogenome of *H. lacustris*, which is inflated by the accumulation of repeats highly similar to those found in the plastome of the same species [11], provided the first evidence that error-prone repair of double-strand breaks leading to repeat proliferation may also occur in chlamydomonadalean mitochondria. Recent study reported the presence of highly similar repeats in the mitogenome of another chlamydomonadalean alga, *Stephanosphaera pluvialis*, and proposed horizontal gene transfer between the *H. lacustris* and *S. pluvialis* lineages as a possible explanation for the sharing of similar mitochondrial repeats by the two organisms [60]. Our characterisation of the *L. pallida* mitogenome, which is also repeat-rich and larger than any chlamydomonadalean algal mitogenome sequenced to date except that from *H. lacustris*, revealed that mitogenome inflation may be more common in Chlamydomonadales. However, in contrast to *H. lacustris*, the GC content as well as the repeats in the two organellar genomes of *L. pallida* differ significantly (Additional file 3: Table S2), so the evolutionary path leading to the parallel inflation of both genomes in this lineage may have been completely different from the one manifested in *H. lacustris*. Strikingly, the specific nature of the mitochondrial repeats in *L. pallida* entails the high abundance of PQS in the mitogenome. G-quadruplexes are increasingly recognised as regulatory structures [61], and they can also form in the mitogenomes, although their role in mtDNA has yet to be elucidated [62]. However, the PQS abundance in the *L. pallida* mitogenome is truly extreme and comparable only with the situation in the mitogenome of the lycophyte *S. moellendorffii* (Additional file 3: Table S2). Both species are thus interesting candidates for studying the role of G-quadruplexes in mitochondrial DNA.

## Conclusions

Our study indicates that continued sampling of microbial eukaryotes is critical for further progress in our knowledge of the phylogenetic diversity of life, and for forming a better understanding of the general principles governing the evolution of organellar genomes. The specific factors contributing to the propensity of chlamydomonadalean organellar genomes to accumulate repetitive sequences, reaching an extreme in *L. pallida*, remain unknown and may not be easy to define. However, future research on *Leontynka*, including characterisation of the organellar genomes of *L. elongata*, may generate additional insights into the molecular mechanisms and evolutionary forces shaping the organellar genomes in this group. It will also be important to perform a detailed comparative analysis of the molecular machinery responsible for genome replication and maintenance in Chlamydomonadales and other green algae. The transcriptome assembly reported here for *L. pallida* will be instrumental not only in this enterprise, but will also serve as a resource for exploring the full range of physiological roles of the plastid in the *Leontynka* lineage, and may help to further clarify the phylogenetic position of *Leontynka* within Chlamydomonadales. We posit that *Leontynka* may become an important model system for analysing the evolutionary and functional aspects of photosynthesis loss in eukaryotes with primary plastids.

## Methods

### Isolation, cultivation, and basic characterisation of new protist strains

The two strains AMAZONIE and MBURUCU were obtained from freshwater hypoxic sediment samples collected in Peru and Argentina, respectively. The strains were cultivated and morphologically characterised by light and transmission electron microscopy, using routine methods. Basic molecular characterisation was achieved by determining partial sequences of the rDNA operon. Further details are provided in Additional file 2: Methods S1-S3.

### Organellar genome and nuclear transcriptome sequencing

Bacterial contamination in the AMAZONIE culture was minimised by filtration, and DNA and RNA were extracted using standard protocols detailed in Additional file 2: Methods S4. Nanopore sequencing was performed using 4 μg of genomic DNA. The DNA was sheared at 20 kbp using Covaris g-TUBE (Covaris) according to the manufacturer’s protocol. After shearing, two libraries were prepared using a Ligation Sequencing Kit from Oxford Nanopore Technologies (SQK-LSK108). The prepared library was loaded onto a R9.4.1 Spot-On Flow cell (FLO-MIN106). Sequencing was performed on a MinION Mk1B machine for 48 hours using the MinKNOW 2.0 software. Basecalling was performed using Guppy 3.0.3 with the Flip-flop algorithm. Illumina sequencing of the genomic DNA was performed using 1 μg of genomic DNA with Illumina HiSeq 2000 (2×150bp) paired-end technology, with libraries prepared using TruSeq DNA PCR-Free (Illumina, San Diego, CA), at Macrogen Inc. (Seoul, South Korea). The transcriptome was sequenced using HiSeq 2000 (2×100bp) paired-end technology, with libraries prepared using the TruSeq RNA sample prep kit v2 (Illumina, San Diego, CA), at Macrogen Inc. (Seoul, South Korea).

### Organellar genome and nuclear transcriptome assembly

Raw Illumina sequencing reads were trimmed with Trimmomatic v0.32 [63]. Initial assembly of the Oxford Nanopore data was performed using Canu v1.7 with the corMaxEvidenceErate set to 0.15 [64]. After assembly, the plastome-derived contigs were identified using BLAST [65], with the *Chlamydomonas reinhardtii* plastome as a query. Nine putative plastid genome sequences were selected and polished using the raw nanopore reads with Nanopolish [66], followed by polishing with Illumina reads with Pilon v1.22 [67]. The Illumina reads were re-mapped onto polished contigs and the mapped reads were then extracted and used as an input for Unicycler v0.4.8 [68] together with the nanopore reads. Unicycler generated a single circular contig of 362,307 bp. For the mitogenome, a single linear contig was identified in the Canu assembly with BLAST using standard mitochondrial genes as queries; the contig sequence was polished using the same method as described above, but it remained linear after a subsequent Unicycler run. However, direct inspection of the contig revealed highly similar regions (about 5,600 bp in length) at both termini. The terminal regions were further polished by mapping of Illumina genomic reads using BWA [69] and SAMtools [70], followed by manual inspection in Tablet [71], which increased the sequence similarity of the termini to 97.7% (along the region of 5,771 bp).

Illumina genomic reads were also assembled separately with the SPAdes Genome assembler v3.10.1 [72], and subsequently used for cleaning the transcriptomic data as follows. Contaminant bacterial contigs >400,000 bp that were identified with BLAST in the SPAdes (16 contigs) and Canu assemblies (11 contigs), together with published genome assemblies of close relatives of bacteria identified in the AMAZONIE culture (*Curvibacter lanceolatus* ATCC 14669, *Bacteroides luti* strain DSM 26991, and *Paludibacter jiangxiensis* strain NM7), were used for RNA-seq read mapping (Hisat2 2.1.0; [73]) to identify and remove bacterial transcriptomic reads that survived the filtration of the culture and polyA selection. This procedure removed ~4% of the reads. Cleaned reads were used for transcriptome assembly with rnaSPAdes v3.13.0 using a k-mer size of 55 bp [74].

### Annotation of organellar genomes and other sequence analyses

Initial annotation of both the plastid and mitochondrial genomes of the strain AMAZONIE were obtained using MFannot (http://megasun.bch.umontreal.ca/cgi-bin/dev_mfa/mfannotInterface.pl). The program output was carefully checked manually, primarily by relying on BLAST searches, to find possible missed genes, to validate or correct the assessment of the initiation codons, to fix the delimitation of introns, and to ensure that all genes were properly named. ORFs encoding proteins shorter than 150 amino acid residues and lacking discernible homologs (as assessed by HHpred [75]) and ORFs consisting mostly of sequence repeats were omitted from the annotation. tRNA genes were identified and annotated by using the bacterial mode of tRNAscan-SE 2.0 [76]. The organellar genome maps were visualised by using OGDRAW v1.3.1 [77]. The distribution of repeats within the organellar genomes and a comparison of repeats between the organellar genomes of *L. pallida* and other selected chlamydomonadaleans were analysed using the dottup program from the EMBOSS package (http://www.bioinformatics.nl/cgi-bin/emboss/dottup). Detailed analyses of imperfect palindromes and G-quadruplexes were performed using the Palindrome analyzer [78] and the G4hunter web-based server [79]. The Palindrome analyzer was used to search for motifs 8-100 bp in length, with spacers of 0-10 bp, and a maximum of one mismatch in the palindrome. The G4hunter web-based server was used with the default settings, i.e., window=25 and threshold=1.2.

RNA-seq reads were mapped onto the plastome sequence using Hisat2 and the result inspected in Tablet in an attempt to check if repeats in intragenic regions could be spliced out as non-standard introns. To understand the position of amino acid stretches encoded by the characteristic repeats that have invaded the coding sequence of the *ftsH* gene, the tertiary structure of the encoded protein was predicted by homology modelling using the Phyre2 program (http://www.sbg.bio.ic.ac.uk/phyre2/html/page.cgi?id=index; [80]). The secondary structure of the ITS2 region was modelled manually according to the consensus secondary ITS2 structure of two green algae [81], visualised by the VARNA software [82], and manually edited in a graphical editor. Homologs of nucleus-encoded plastidial proteins of specific interest were searched for in the *L. pallida* transcriptome assembly by using TBLASTN and the respective protein sequences from *Arabidopsis thaliana* or *C. reinhardtii* (selected based on information from the literature or keyword database searches). Significant hits (E-value ≤1e-5) were evaluated by BLASTX searches against the NCBI non-redundant protein sequence database to filter out bacterial contaminants and sequences corresponding to non-orthologous members of broader protein families. Subcellular localization (for complete sequences only) was assessed by using TargetP-2.0 (https://services.healthtech.dtu.dk/service.php?TargetP-2.0; [83]) and PredAlgo (http://lobosphaera.ibpc.fr/cgi-bin/predalgodb2.perl?page=main; [84]).

### Phylogenetic analyses

Multiple sequence alignment of the 18S rRNA gene from a total of 201 chlorophyte OTUs was computed using MAFFT v7 [85] and trimmed manually. The 18S rRNA sequence from *Polytoma oviforme* available in GenBank (U22936.1) was proposed to be chimeric [12], but given the relevance of this organism for our analysis, we included it, masking the regions putatively derived from a different source by strings of N. Maximum likelihood tree inference was performed using IQ-TREE multicore v1.6.12 [86] using the TIM2+F+I+G4 model with 100 non-parametric bootstrap replicates. For multigene analysis, alignments of conserved plastome- and mitogenome-encoded proteins used previously [87,88] were updated by adding the respective homologs from *L. pallida* and additional relevant chlorophycean taxa not represented in the initial datasets. Sequences representing only distantly related outgroup taxa, namely the OCC clade of Chlorophyceae (for the plastome-based dataset) and Ulvophyceae (for the mitogenome-based dataset) were removed from the analyses to keep the size of the dataset easier to analyse with a complex substitution model. For the final matrices, a subset of 24 plastid-encoded and seven mitogenome-encoded proteins (all with their *L. pallida* representative) were used. Multiple alignments of the homologous amino acid sequences were built using MAFFT v7.407 with the L-INS-i algorithm [89], and were manually trimmed to exclude unreliably aligned regions. The final concatenated matrix comprised 5,020 amino acid residues. The tree was built using PhyloBayes v4.1 [90] using the CAT+GTR model of sequence evolution, with two independent chains that converged at 15,298 generations, with the largest discrepancy in posterior probabilities (PPs) (maxdiff) being 0.0535238 (at burn-in of 20%). The maximum likelihood (ML) tree was inferred with IQ-TREE multicore v1.6.12 using the LG+C60+F+G4 substitution model. Statistical support was assessed with 100 IQ-TREE non-parametric bootstraps with correction, and additionally with PhyloBayes posterior probabilities for the plastome-based dataset.

## Supplementary information

**Supplementary information** accompanies this paper at XXXXXXX.

**Additional file 1: Supplementary Figures. Fig. S1.** Maximum likelihood phylogenetic tree ((IQ-TREE, TIM2+F+I+G4 substitution model) of 18S rRNA gene sequences from Chlorophyceae. **Fig. S2.** Predicted secondary structure of the ITS2 region of *Leontynka pallida*, with differences in the corresponding region of *Leontynka elongata* mapped onto it. **Fig. S3.** Maximum likelihood phylogenetic tree of Chlorophyceae, including *Leontynka pallida*, inferred from a concatenated set of seven conserved mitogenome-encoded proteins (2,608 amino acid positions). **Fig. S4.** Light micrographs of *Leontynka pallida*. **Fig. S5.** Light micrographs of *Leontynka elongata*. **Fig. S6.** Ultrastructure of *Leontynka elongata* (a–f) and *Leontynka pallida* (g–i). **Fig. S7.** Occurrence of the “variant 8” repeat (see Fig. 4) in the FtsH protein of Leontynka pallida mapped onto its predicted structure. **Fig. S8.** Alignment of the highly similar terminal regions of the originally assembled linear mitogenome contig. **Fig. S9.** Occurrence of the “variant 8” repeat (translated in reading frame +0 as KDKPANLTS and −0 as KEVSFAGLSL; both boxed in colour) in a variable region of protein sequence of the ribosomal protein Rps8 from *Leontynka pallida* (full protein alignment together with representatives of other chlamydomonadalean algae).

**Additional file 2: Supplementary Notes and supplementary Methods. Note S1.** Taxonomic descriptions. **Note S2.** Further details on the morphology and ultrastructure of *Leontynka* spp. **Note S3.** Further details on various kinds of repeats in the plastome of *L. pallida*. **Note S4.** Further details on the repeat insertions in the *L. pallida* plastid coding sequences. **Note S5.** Differential diagnosis of *Leontynka* spp. with regard to previously described colourless chlamydomonadalean taxa. **Methods S1.** Isolation and cultivation of strains. **Methods S2.** Light and transmission electron microscopy. **Methods S3.** Amplification and sequencing of 18S and ITS rDNA regions. **Methods S4.** DNA and RNA isolation.

**Additional file 3: Supplementary Tables. Table S1.** Nuclear transcripts from *Leontynka pallida* specifically discussed in the paper. **Table S2.** Comparison of GC content, number of imperfect palindromes, and potential quadruplex-forming sequences in selected organellar genomes. **Table S3.** Strong codon usage bias in the mitochondrial genome of *Leontynka pallida*. **Table S4.** Relative frequency of amino acids with GC-rich codons (G, A, R, P) in proteins encoded by different mitogenomes. **Table S5.** Relative frequency of codons in plastid genes of *Leontynka pallida*. **Table S6.** Relative frequency of amino acids in proteins encoded by the plastome of *Leontynka pallida*. **Table S7.** The most abundant imperfect palindrome in the *Leontynka pallida* plastome that is missing in the exons.

## Declarations

### Ethics approval and consent to participate

Not applicable.

### Consent for publication

Not applicable.

### Competing interests

The authors declare no competing interests.

### Availability of data and materials

All data needed to evaluate the conclusions in the paper are present in the paper, the Supplementary Information files and/or publicly available repositories. rDNA and organellar genome sequences determined in this study are available from GenBank with the following accession numbers: OM501587.1 [91] and OM501588.1 [92], OM479424.1 [93], and OM479425.1 [94]. Genomic and transcriptomic reads were deposited at NCBI as the BioProject PRJNA799256 [95]. The transcriptome assembly from *L. pallida* as well as single-gene datasets and concatenated protein sequence matrices used for phylogenomic analyses are available from Figshare [96–98]. The cultures of *Leontynka* spp. investigated in this study are available upon request. The type material for both newly described species was deposited at the National Museum, Prague, Czech Republic; inventory number P6E 5164-5165.

## Abbreviations

ATP: Adenosine triphosphate
CBCs: Compensatory base changes
DNA: Deoxyribonucleic acid
ITS: Internal transcribed spacer
IP: imperfect palindrome
ML: Maximum likelihood
MYA: Million year ago
ORF: Open reading frame
PQS: potential quadruplex-forming sequences
RNA: Ribonucleic acid
rRNA: Ribosomal ribonucleic acid
SRP: Signal recognition particle
TAT: Twin-arginine protein translocase
tRNA: Transfer ribonucleic acid

## Funding

This work was supported by the Czech Science Foundation project 17-21409S (to ME) and the project “CePaViP”, supported by the European Regional Development Fund, within the Operational programme for Research, Development and Education (CZ.02.1.01/0.0/0.0/16_019/0000759). TP was also supported by Charles University (UNCE 204069).

## Authors’ contributions

TP, DB, IČ and ME conceived the original research plans; TP, SCT, KZ, EZ, and KJ obtained nucleic acids for sequencing; SCT and TP obtained Oxford Nanopore data and generated organellar genome assemblies; EZ and IČ isolated the strains; TP and DB carried out the morphological characterisation of the strains; DB and NY obtained the TEM data; MS assembled the transcriptome; TP, DB, and TŠ carried out phylogenetic analyses; TP, IČ and ME analysed and annotated the organellar genome sequences; IČ and ME supervised the work of junior researchers and obtained funding; TP, DB and ME drafted the manuscript; all authors contributed to the text; all authors read and approved the final manuscript; ME agreed to serve as the author responsible for contact and ensuring communication.

## Acknowledgements

We are thankful to Karolina Fučíková for providing alignments of chlorophycaean plastid proteins, Monique Turmel and Claude Lemieux for providing alignments of chlorophyceaen mitochondrial proteins, Petr Janšta for collecting the sample MBURUCU, and the anonymous reviewers for their useful comments on the previous versions of the manuscript. TP thanks IT4Innovations National Super Computing Center, VSB – Technical University of Ostrava, Czech Republic (project #Open-22-48) for providing computational resources.

## Notes

### Competing Interest Statement

The authors have declared no competing interest.

