## Additional file 1: Supplementary Figures for "A new lineage of non-photosynthetic green algae with extreme organellar genomes"

**This file contains supplementary Figs S1 to S9.**

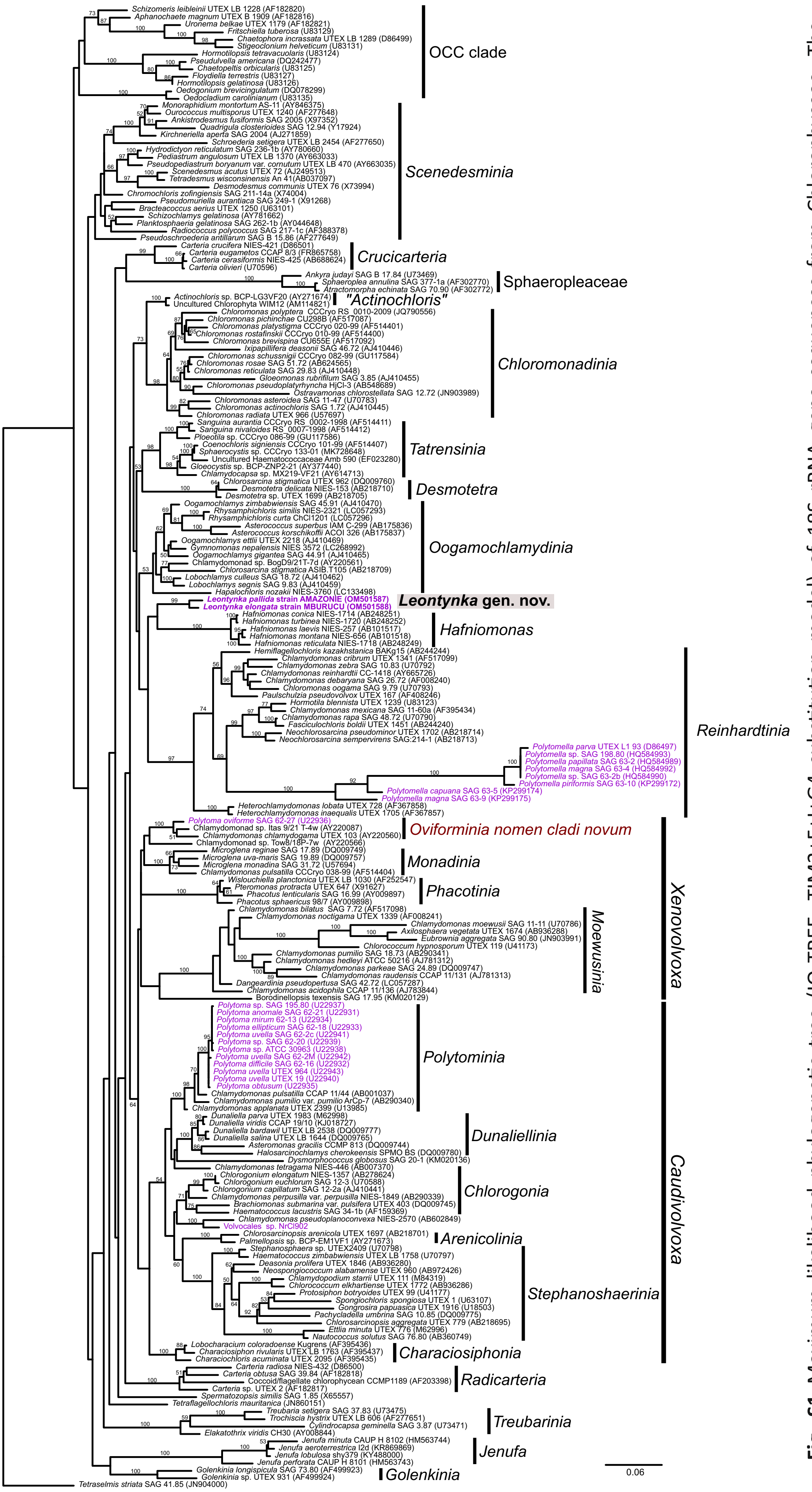


**Fig. S1** Maximum likelihood phylogenetic tree (IQ-TREE, TIM2+F+I+G4 substitution model) of 18S rRNA gene sequences from Chlorophyceae. The chlorodendrophyceaen *Tetraselmis striata* is used as an outgroup. Bootstrap support values are shown when ≥50. Previously demarcated main clades [12] are annotated in the tree together with the newly designated clade *"Oviforminia"*. Sequences from non-photosynthetic taxa are in colour. The new genus *Leontynka* (highlighted) forms a novel clade without apparent specific affinities to other particular lineages of Chlamydomonadales. The clade *"Actinochloris"* is labelled provisionally, as the *bona fide Actinochloris* genus belongs elsewhere.

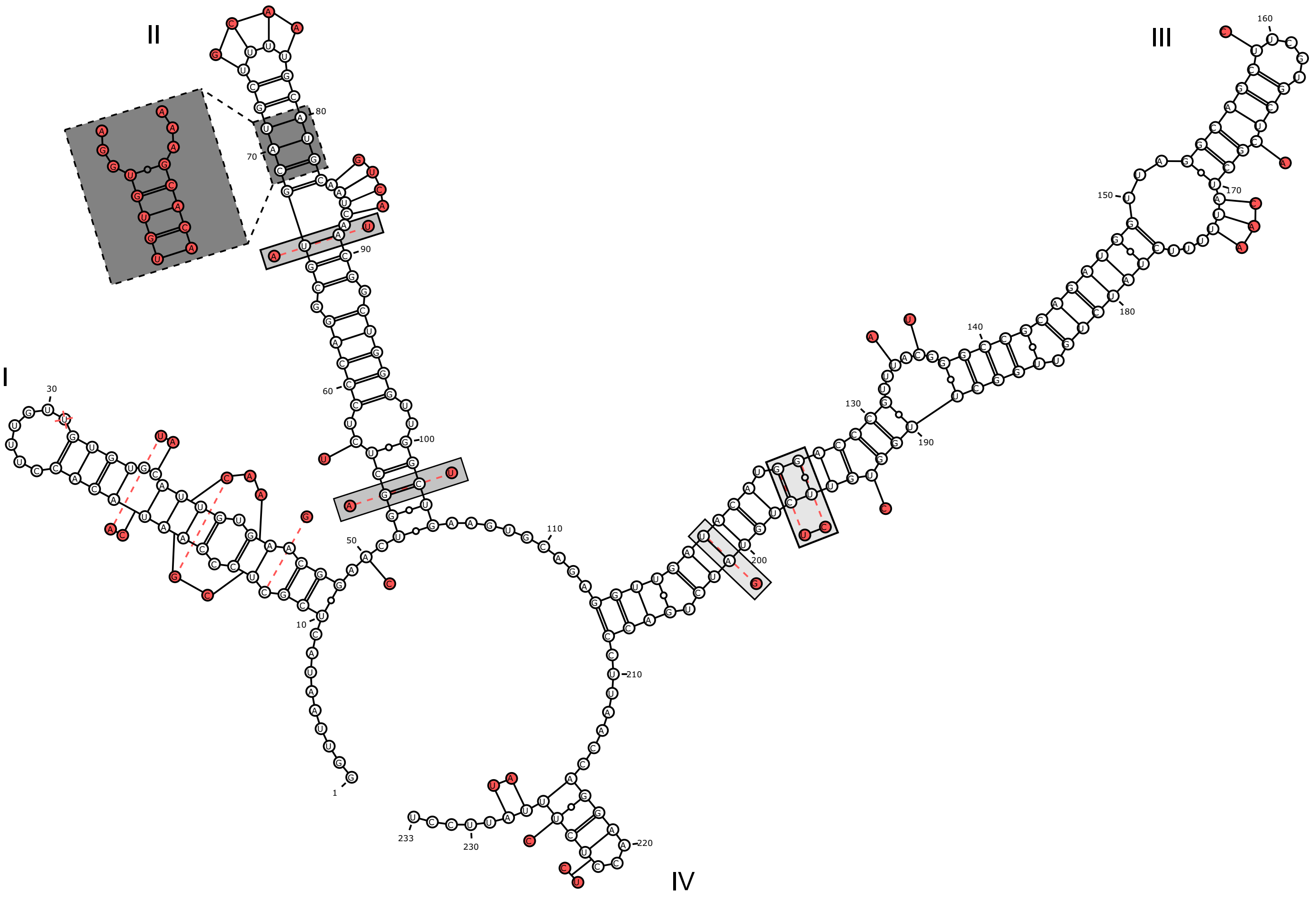


**Fig. S2** Predicted secondary structure of the ITS2 region of *Leontynka pallida*, with differences in the corresponding region of *Leontynka elongata* mapped onto it. Classical compensatory base changes in helix II are highlighted by a light grey backround, and a region more substantially differing between the two species is highlighted by a dark grey background.

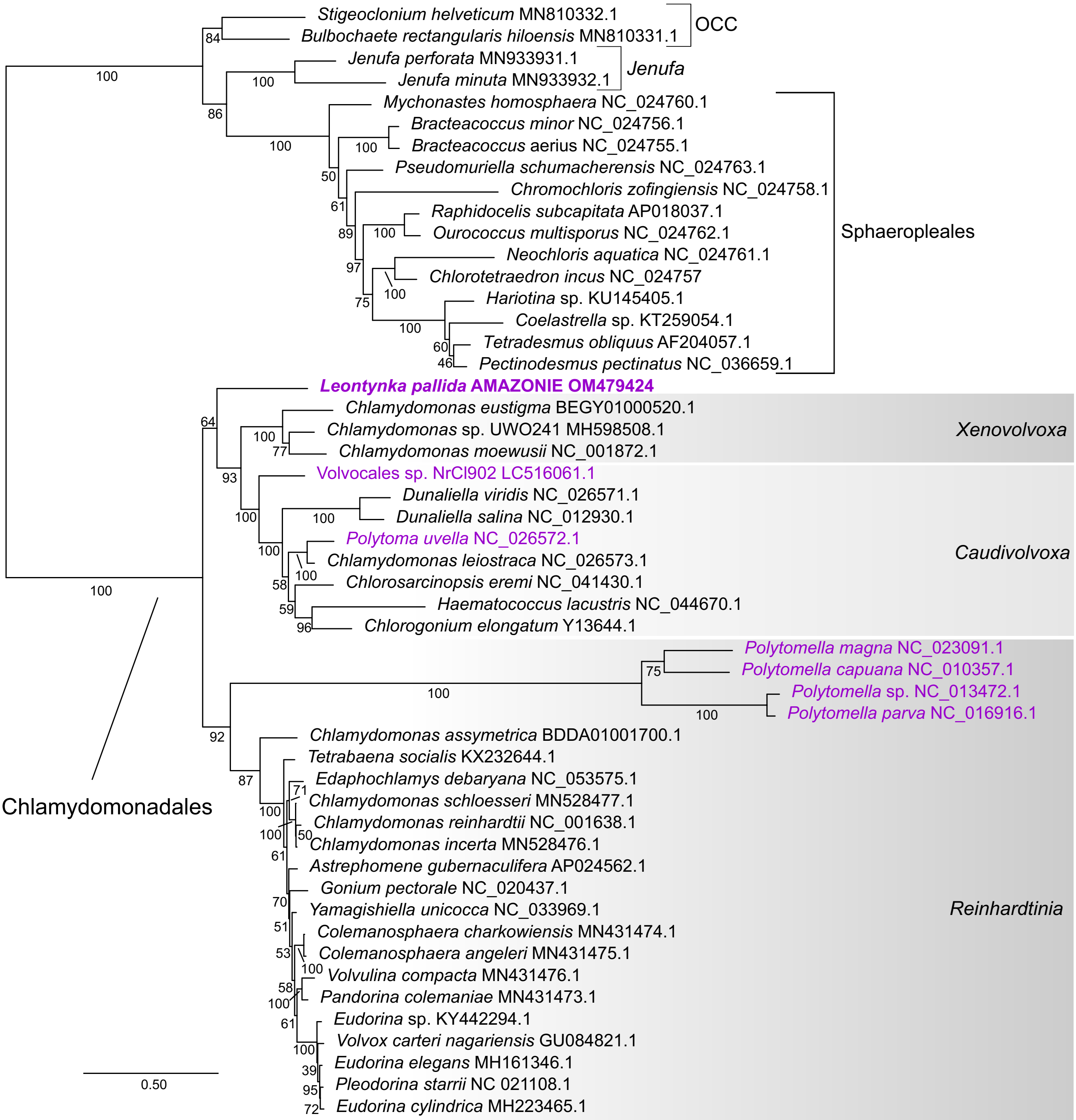


**Fig. S3** Maximum likelihood phylogenetic tree of Chlorophyceae, including *Leontynka pallida*, inferred from a concatenated set of seven conserved mitogenome-encoded proteins (2,608 amino acid positions). The tree topology was inferred using maximum likelihood analysis (IQ-TREE, LG+C60+F+G4 substitution model, 100 non-parametric bootstrap replicates). For easy display, the tree is arbitrarily rooted between Chlamydomonadales and other Chlorophyceae included in the analysis.

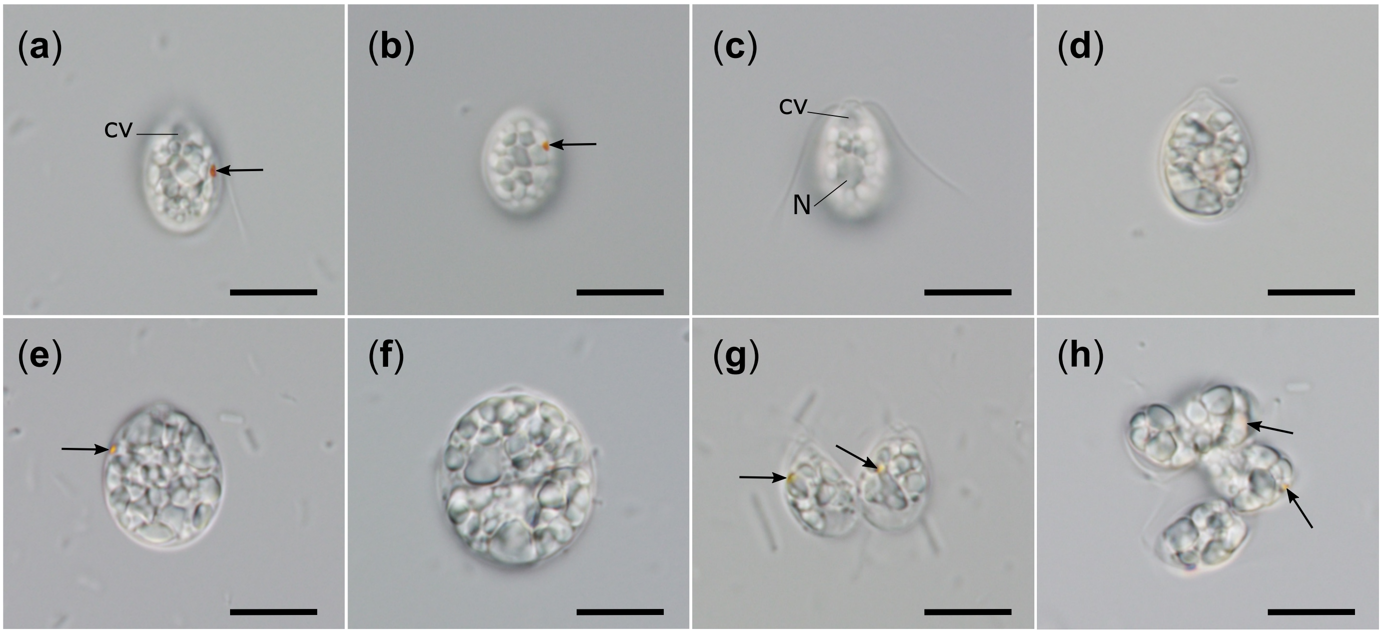


**Fig. S4** Light micrographs of *Leontynka pallida*. Scale bars = 10 µm.
Abbreviations: arrows – eyespot; cv – contractile vacuole; N – nucleus.

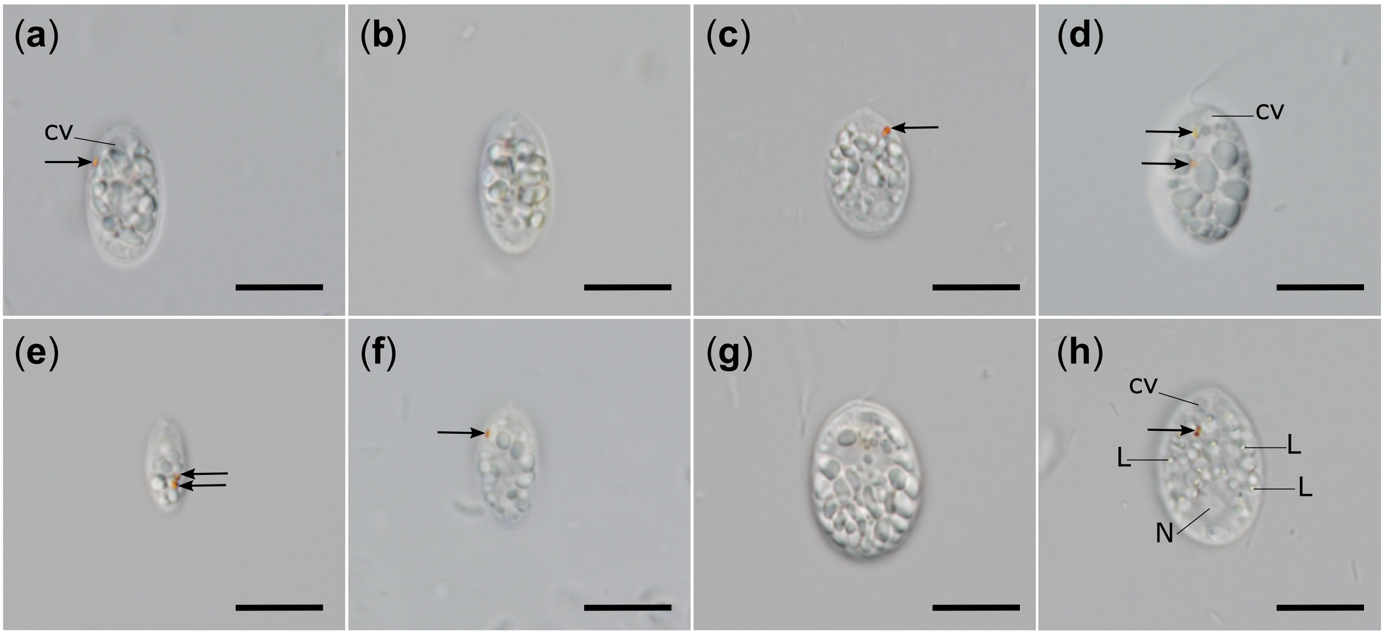


**Fig. S5** Light micrographs of *Leontynka elongata*. Scale bars = 10 µm.
Abbreviations: arrows – eyespot; cv – contractile vacuole; L – lipid droplet; N – nucleus.

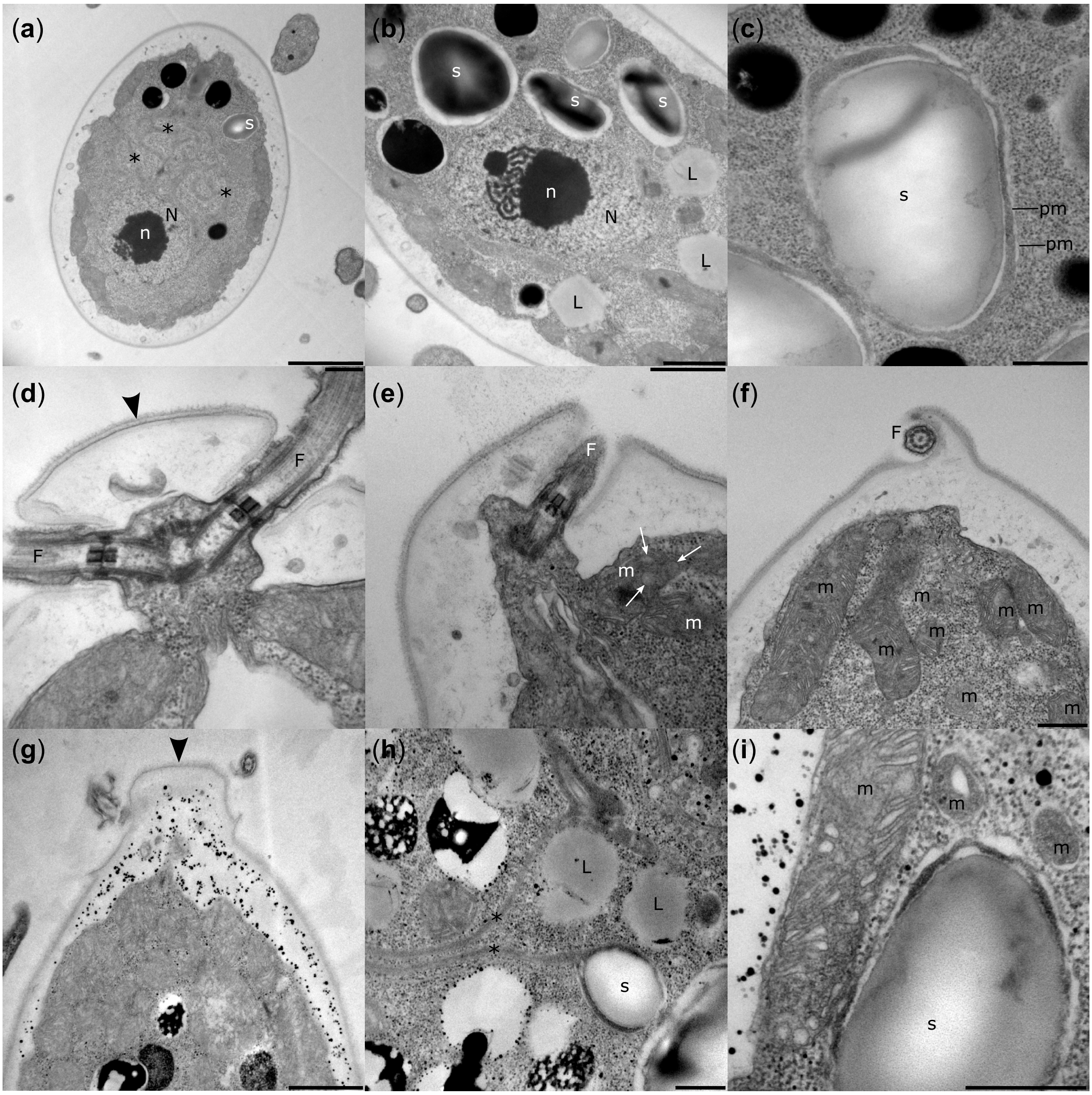


**Fig. S6** Ultrastructure of *Leontynka elongata* (a–f) and *Leontynka pallida* (g–i). (a) Cell with a slightly posterior nucleus and highly convoluted leucoplast. (b) Cell with a central nucleus and multiple starch blocks. (c) Two membranes surround the plastid. (d) Prominent keel-shaped papilla and two flagella. (e) Cross section through mitochondria with discoidal cristae. (f) Longitudinal section through mitochondria with discoidal cristae. (g) Ovoid cell with a keel-shaped papilla. (h) Presence of lipid droplets in an older cell. (i) Mitochondria containing putative tubulo-vesicular cristae (longitudinal section through the organelle). Abbreviations: F – flagellum; L – lipid droplet; m – mitochondrion; N – nucleus; n – nucleolus; pm – plastid membrane; s – starch. Asterisks mark “bridges” between plastid compartments; black arrowheads indicate papillae; white arrows indicate discoidal cristae. Scale bars: a = 2 µm; b, g = 1 µm; c, f, h, i = 0.5 µm d, e = 0.2 µm.

### Phyre2

Date: Fri Nov 13 17:28:33 GMT 2020

Confidence Key
High(9) — Low (0)
? Disordered ( 30%)
Alpha helix ( 54%)
Beta strand ( 11%)
TM helix ( 2%)

Secondary structure and disorder prediction

FtsH - *Leontynka pallida*

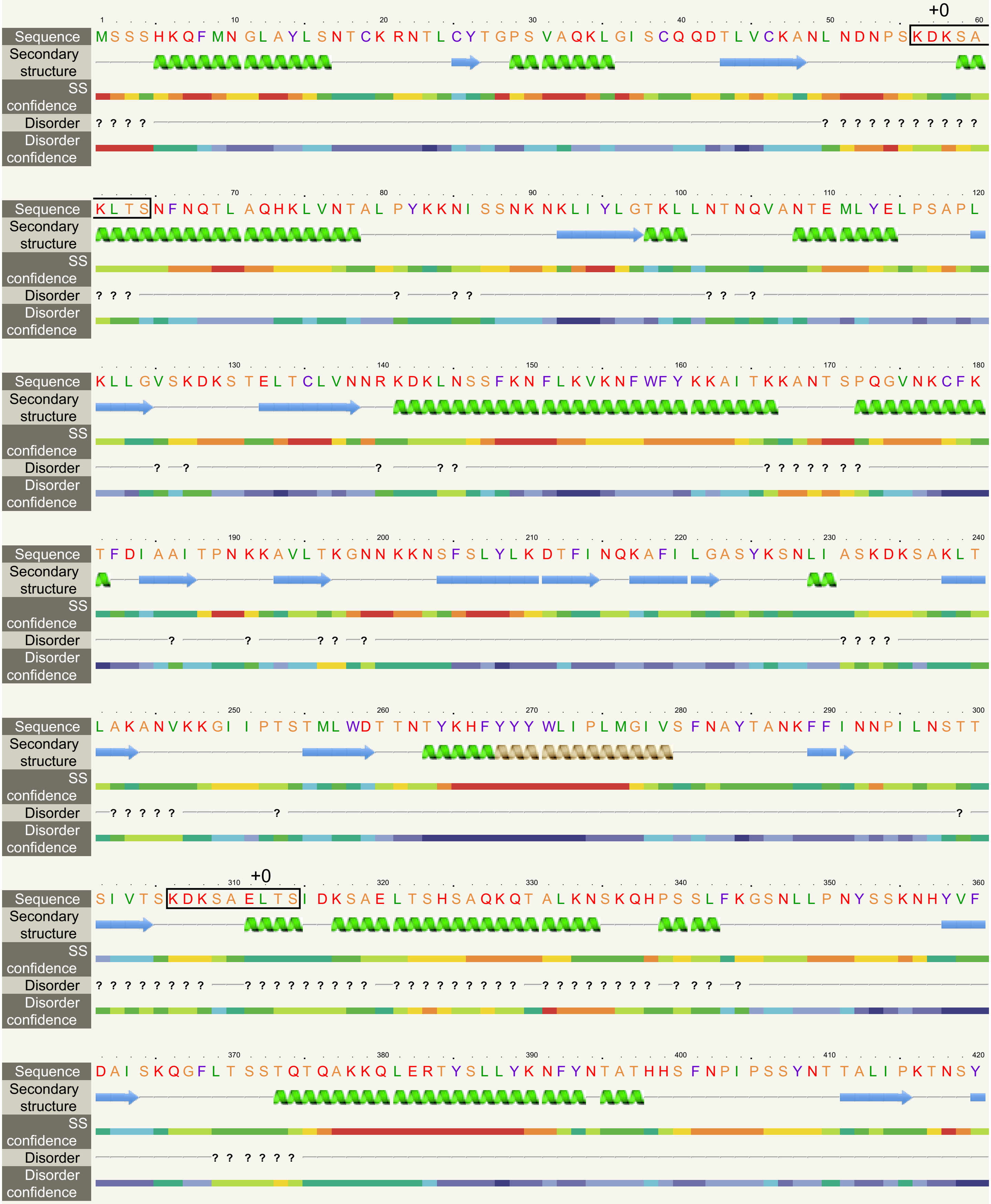

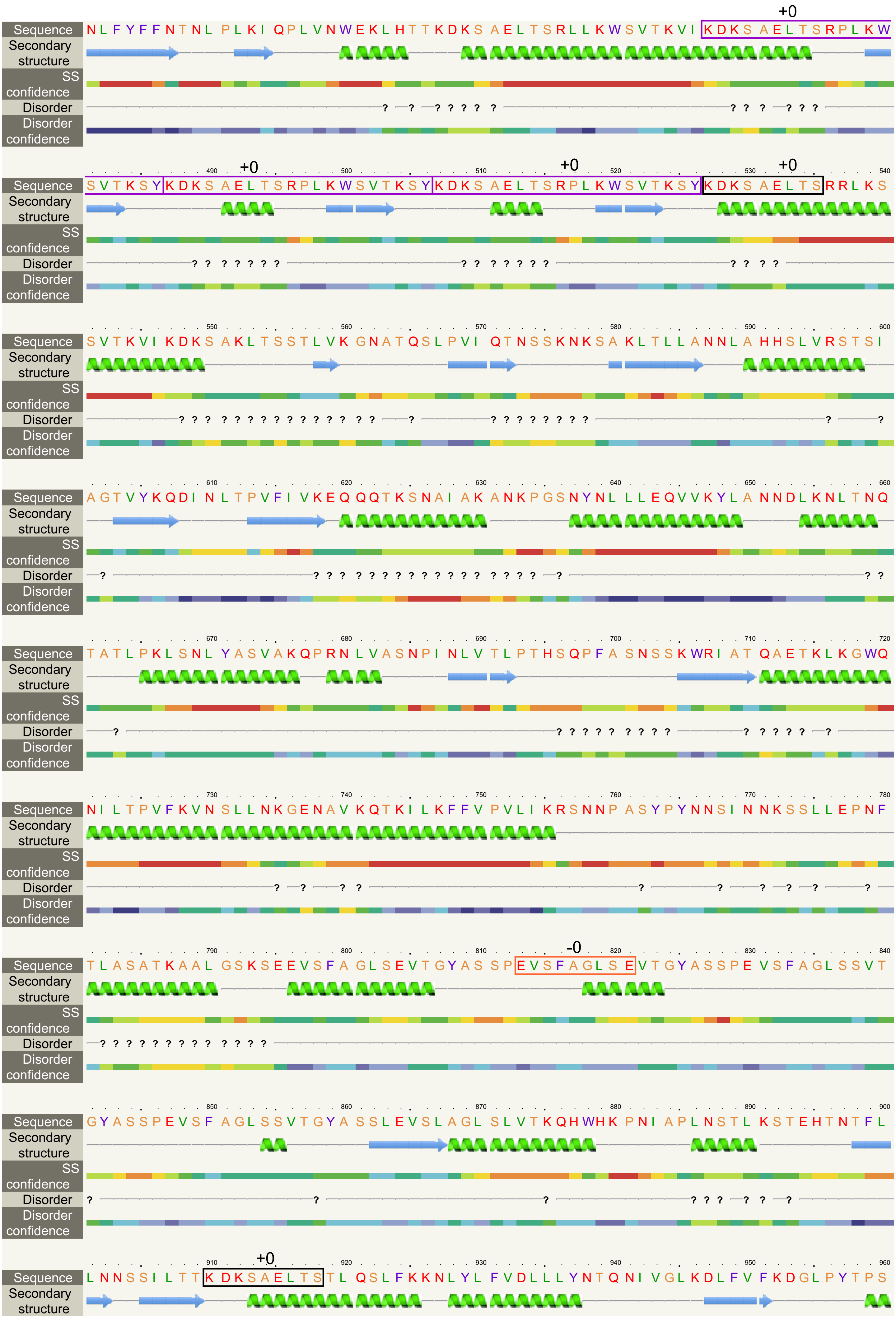
Sequence
Secondary structure
SS confidence
Disorder
Disorder confidence
+0
NLFYFFNTNL PLKI QPLVNWEKLHTTKDKS AELTSRLLKWSVTKVI KDKS AELTSRPLKW
490 500 510 520 530 540
+0 +0 +0
SVTKSY KDKS AELTSRPLKWSVTKSY KDKS AELTSRPLKWSVTKSY KDKS AELTS RRLKS
550 560 570 580 590 600
SVTKVI KDKS AKLTSSTLVK GNATQSLPVI QTNSSKNKSA KLTLLANNLA HHSLVRSTSI
610 620 630 640 650 660
AGTVYKQDI NLTPVFI VKEQ QQTKSNAI AK ANKPGSNYNL LLEQVVKYLA NNDLKNLTNQ
670 680 690 700 710 720
TATLPKLSNL YASVAKQPRN LVASNPI NLV TLPTHSQPFA SNSSKWRI AT QAETKLKGWQ
730 740 750 760 770 780
NI LTPVFKVN SLLNKGENAV KQTKI LKFFV PVLI KRSNNP ASYPYNNSI NNKSSLLEPNF
790 800 810 820 830 840
-0
TLASATKAAL GSKSEEVSFA GLSEVTGYAS SPEVSFAGLS EVTGYASSPE VSFAGLSSVT
850 860 870 880 890 900
GYASSPEVSF AGLSSVTGYA SSLEVSLAGL SLVTKQHWHK PNI APLNSTL KSTEHTNTFL
910 920 930 940 950 960
+0
LNNSSI LTTK DKSAELTSTL QSLFKKNLYL FVDLLLYNTQ NI VGLKDLFV FKDGLPYTPS

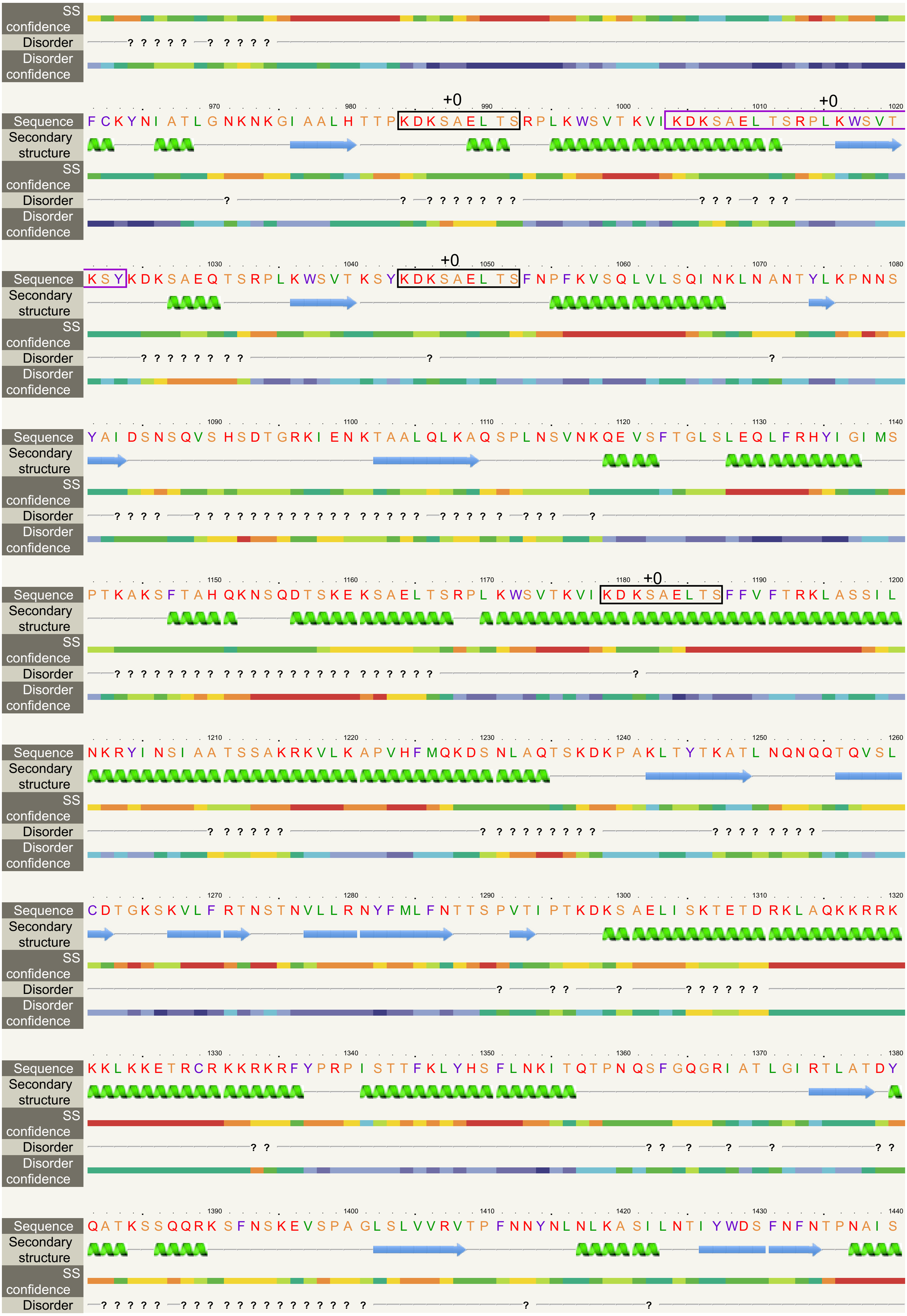

SS confidence
Disorder
Disorder confidence
970 980 +0 990 1000 +0 1010 1020
Sequence FCKYNI ATLGNKNKGI AALHTTPKDKSAELTSRPLKWSVTKVIKDKSAELTSRPLKWSVT
Secondary structure
SS confidence
Disorder
Disorder confidence
1030 1040 +0 1050 1060 1070 1080
Sequence KSYKDKSAEQTSRPLKWSVTKSYKDKSAELTSFNPFKVSQLVLSQINKLNANTYLKPNNS
Secondary structure
SS confidence
Disorder
Disorder confidence
1090 1100 1110 1120 1130 1140
Sequence YAIDSNSQVSHSDTGRKIENKTAALQLKAQSPLNSVNKQEVSFTGLSLEQLFRHYIGIMS
Secondary structure
SS confidence
Disorder
Disorder confidence
1150 1160 1170 1180 +0 1190 1200
Sequence PTKAKSFTAHQKNSQDTSKEKSAELTSRPLKWSVTKVIKDKSAELTSFFVFTRKLASSIL
Secondary structure
SS confidence
Disorder
Disorder confidence
1210 1220 1230 1240 1250 1260
Sequence NKRYINSIAATSSAKRKVLKAPVHFMQKDSNLAQTSKDKPAKLTYTKATLNQNQQTQVSL
Secondary structure
SS confidence
Disorder
Disorder confidence
1270 1280 1290 1300 1310 1320
Sequence CDTGKSKVLFRTNSTNVLLRNYFMLFNTTSPVTIPTKDKSAELISKTETDRKLAQKKRRK
Secondary structure
SS confidence
Disorder
Disorder confidence
1330 1340 1350 1360 1370 1380
Sequence KKLKKETRCRKKRKRFYPRPISTTFKLYHSFLNKITQTPNQSFGQGRIATLGIRTLATDY
Secondary structure
SS confidence
Disorder
Disorder confidence
1390 1400 1410 1420 1430 1440
Sequence QATKSSQQRKSFNSKEVSPAGLSLVVRVTPFNNYNLNLKASILNTIYWDSFNFNTPNAIS
Secondary structure
SS confidence
Disorder

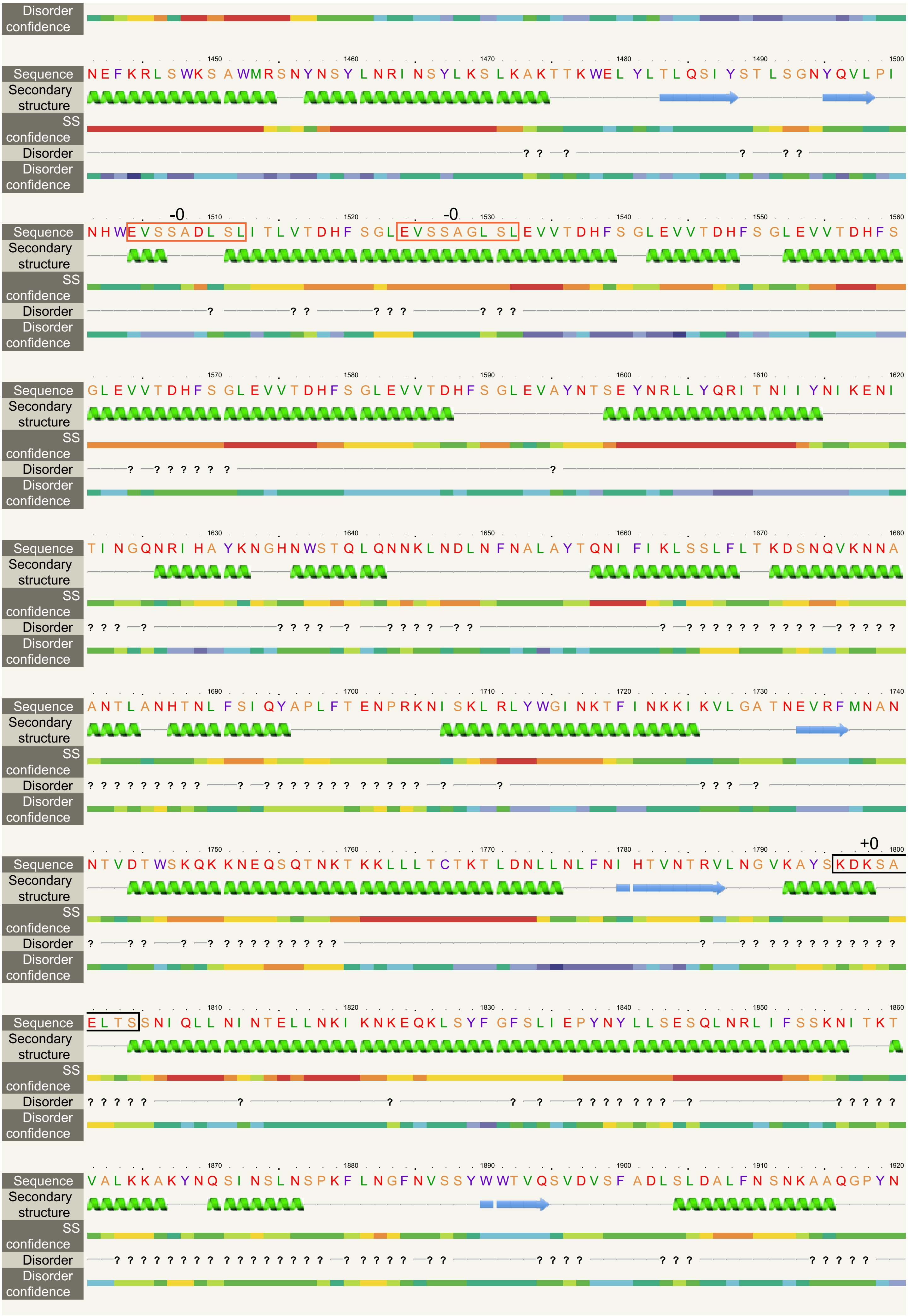
Disorder confidence
Sequence
Secondary structure
SS confidence
Disorder
Disorder confidence
1450 1460 1470 1480 1490 1500
NEFKRLSWKSAWMRSNYNSYLNRINSYLKSLKAKTTKWELYLTLQSIYSTLSGNYQVLPI
-0 -0
1510 1520 1530 1540 1550 1560
NHWEVSSADLSLITLVTDHFSGLEVSSAGLSLEVVTDHFSGLEVVTDHFSGLEVVTDHFS
1570 1580 1590 1600 1610 1620
GLEVVTDHFSGLEVVTDHFSGLEVVTDHFSGLEVAYNTSEYNRLLYQRITNIIYNIKENI
1630 1640 1650 1660 1670 1680
TINGQNRIHAYKNGHNWSTQLQNNKLNDLNFNALAYTQNIFIKLSSLFLTKDSNQVKNNA
1690 1700 1710 1720 1730 1740
ANTLANHTNLFSIQYAPLFTENPRKNISKLRLYWGINKTFINKKIKVLGATNEVRFMNAN
+0
1750 1760 1770 1780 1790 1800
NTVDTWSKQKKNEQSQTNKTKKLLLTCTKTLDNLLNLFNIHTVNTRVLNGVKAYSKDKSA
1810 1820 1830 1840 1850 1860
ELTSSNIQLLNINTELLNKIKNKEQKLSYFGFSLIEPYNYLLSESQLNRLIFSSKNITKT
1870 1880 1890 1900 1910 1920
VALKKAKYNQSINSLNSPKFLNGFNVSSYWWTVQSVDVSFADLSLDALFNSNKAAQGPYN

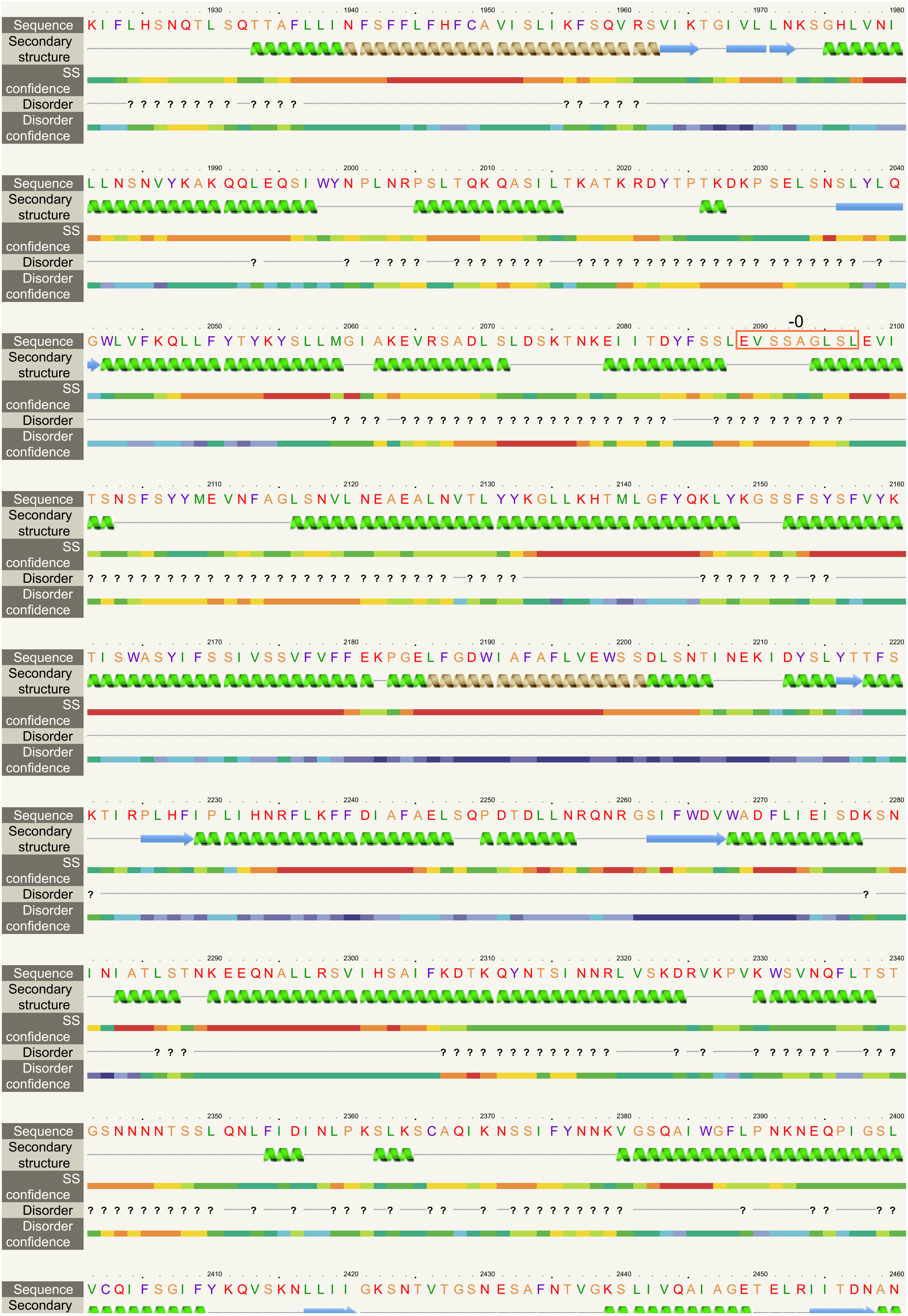

Sequence
Secondary structure
SS confidence
Disorder
Disorder confidence
1930 1940 1950 1960 1970 1980
KIFLHSNQTLSQTTAFLLINFSFFLFHFCAVISLIKFSQVRSVIKTGIVLLNKSGHLVNI
1990 2000 2010 2020 2030 2040
LLNSNVYKAKQQLEQSIWYNPLNRPSLTQKQASILTKATKRDYTPTKDKPSELSNSLYLQ
2050 2060 2070 2080 2090 2100
-0
GWLVFKQLLFYTYKYSLLMGIAKEVRSADLSLDSKTNKEIITDYFSSLEVSSAGLSLEVI
2110 2120 2130 2140 2150 2160
TSNSFSYYMEVNFAGLSNVLNEAEALNVTLYYKGLLKHTMLGFYQKLYKGSSFSYSFVYK
2170 2180 2190 2200 2210 2220
TISWASYIFSSIVSSVFVFFEKPGELFGDWIAFAFLVEWSSDLSNTINEKIDYSLYTTFS
2230 2240 2250 2260 2270 2280
KTIRPLHFIPLIHNRFLKFFDIAFAELSQPDTDLLNRQNRGSIFWDVWADFLIEISDKSN
2290 2300 2310 2320 2330 2340
INIATLSTNKEEQNALLRSVIHSAIFKDTKQYNTSINNRLVSKDRVKPVKWSVNQFLTST
2350 2360 2370 2380 2390 2400
GSNNNNTSSLQNLFIDINLPKSLKSCAQIKNSSIFYNNKVGSQAIWGFLPNKNEQPIGSL
2410 2420 2430 2440 2450 2460
VCQIFSGIFYKQVSKNLLIIGKSNTVTGSNESAFNTVGKSLIVQAIAGETELRIITDNAN

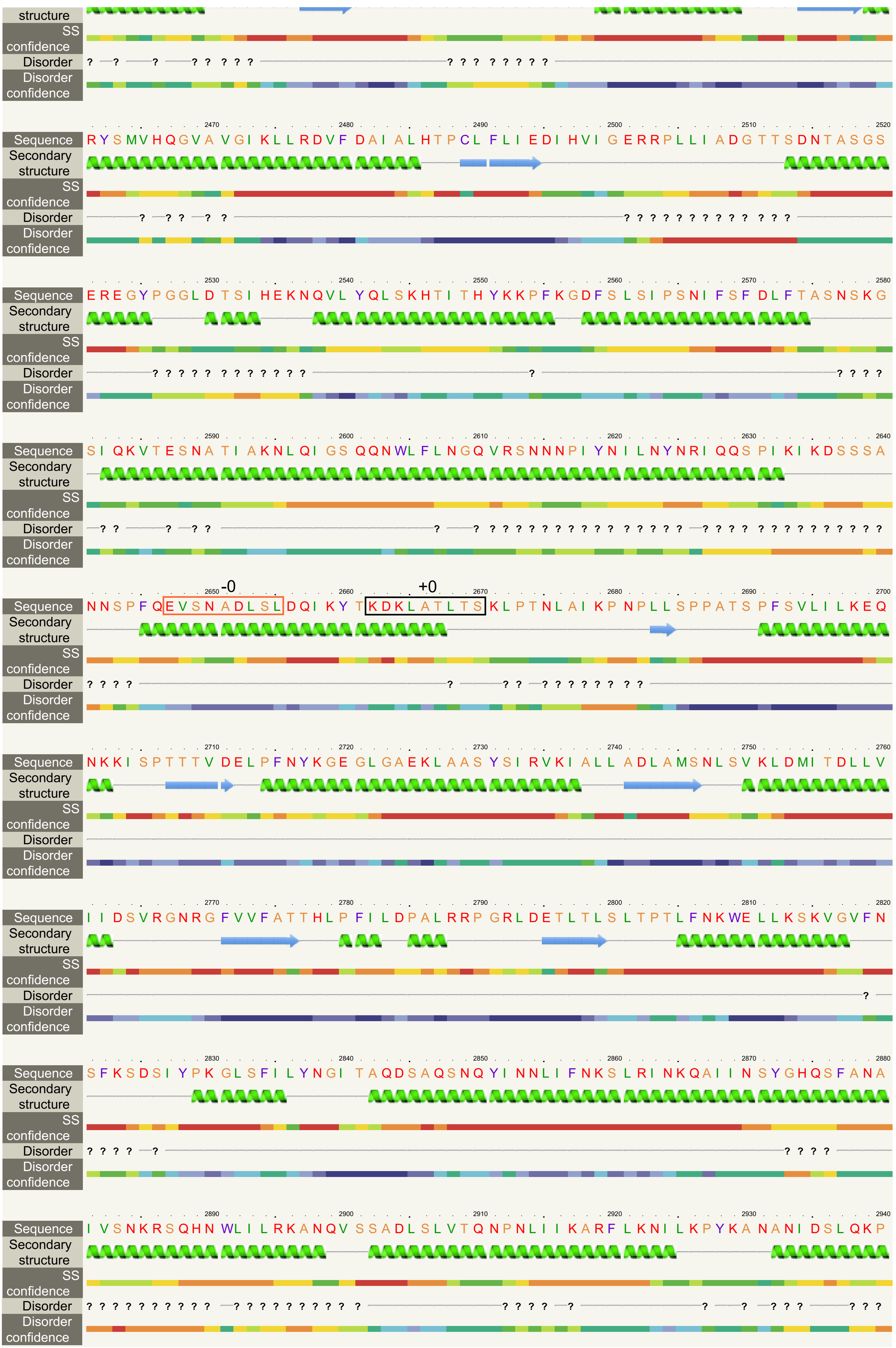

structure
SS confidence
Disorder
Disorder confidence
2470 2480 2490 2500 2510 2520
Sequence RYSMVHQGVAVGIKLLRDVFDAIALHTPCLFLIEDIHVIGERRPLLIADGTTSDNTASGS
Secondary structure
SS confidence
Disorder
Disorder confidence
2530 2540 2550 2560 2570 2580
Sequence EREGYPGGLDTSIHEKNQVLYQLSKHTITHYKKPFKGDFSLSIPSNIFSFDLFTASNSKG
Secondary structure
SS confidence
Disorder
Disorder confidence
2590 2600 2610 2620 2630 2640
Sequence SIQKVTESNATIAKNLQIGSQQNWLFLNGQVRSNNNPIYNILNYNRIQQSPIKIKDSSSA
Secondary structure
SS confidence
Disorder
Disorder confidence
2650 -0 2660 +0 2670 2680 2690 2700
Sequence NNSPFQEVSNADLSLDQIKYTKDKLATLTSKLPTNLAIKPNPLLSPPATSPFSVLILKEQ
Secondary structure
SS confidence
Disorder
Disorder confidence
2710 2720 2730 2740 2750 2760
Sequence NKKISPTTTVDELPFNYKGEGLGAEKLAASYSIRVKIALLADLAMSNLSVKLDMITDLLV
Secondary structure
SS confidence
Disorder
Disorder confidence
2770 2780 2790 2800 2810 2820
Sequence IIDSVRGNRGFVVFATTHLPFILDPALRRPGRLDETLTLSLTPTLFNKWELLKSKVGVFN
Secondary structure
SS confidence
Disorder
Disorder confidence
2830 2840 2850 2860 2870 2880
Sequence SFKSDSIYPKGLSFILYNGITAQDSAQSNQYINNLIFNKSLRINKQAIINSYGHQSFANA
Secondary structure
SS confidence
Disorder
Disorder confidence
2890 2900 2910 2920 2930 2940
Sequence IVSNKRSQHNWLILRKANQVSSADLSLVTQNPNLIIKARFLKNILKPYKANANIDSLQKP
Secondary structure
SS confidence
Disorder
Disorder confidence

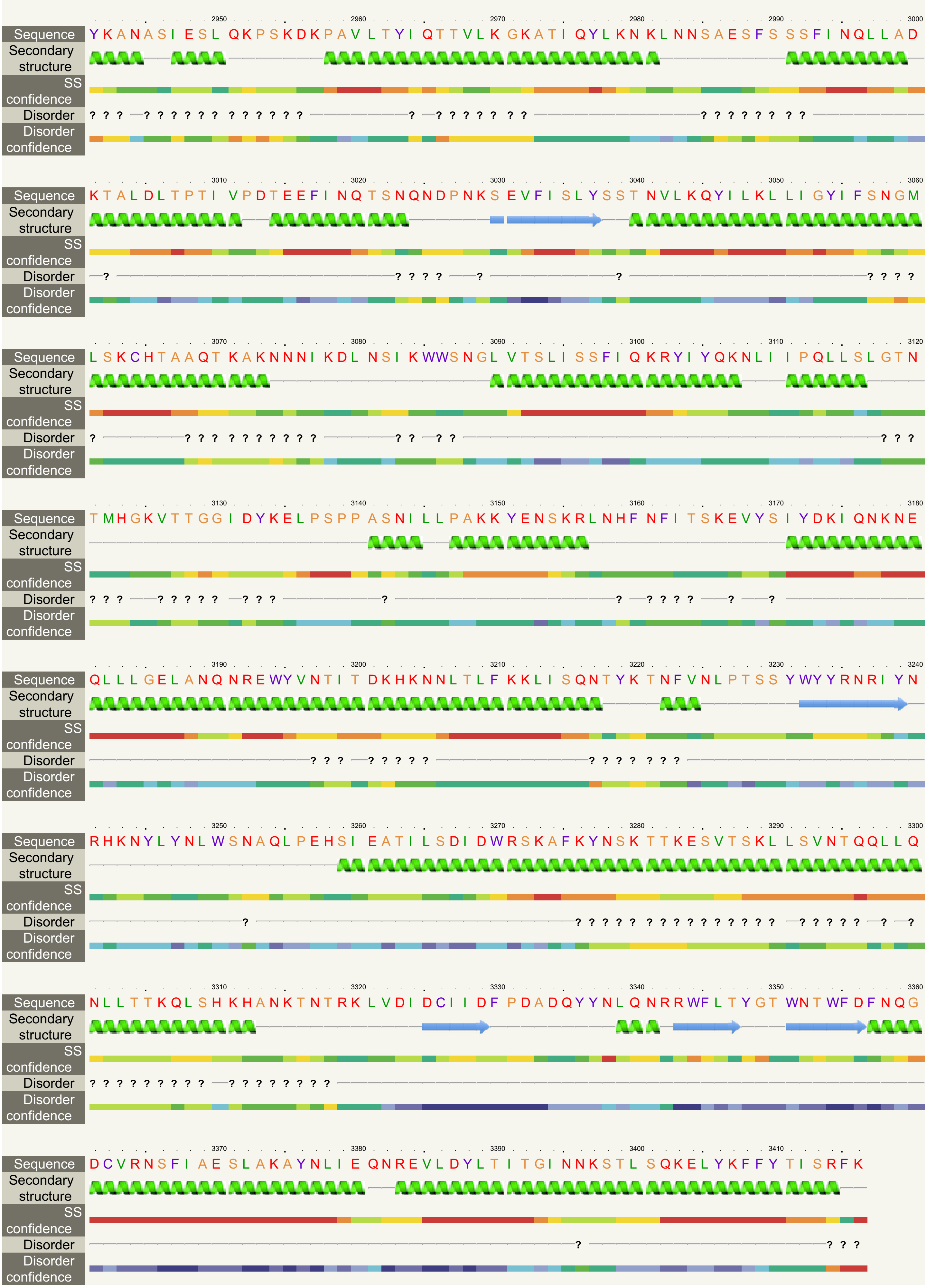

**Fig. S7** Occurrence of the "variant 8" repeat (see Fig. 4) in the FtsH protein of *Leontynka pallida* mapped onto its predicted structure. Protein model was constructed *in silico* using Phyre2. The "variant 8" repeat in RF +0 and -0 as well as a larger repeat containing the "variant 8" repeat in RF +0 are highlighted in coloured boxes.

```
                        10        20        30        40        50        60        70        80        90       100
               ....|....|....|....|....|....|....|....|....|....|....|....|....|....|....|....|....|....|....|....|
realigned-end  AACATGTTCTCTTGTGGCTCGTCCGAGTCGCTCGGCTTCGCCTCGCTCCTCGTCCCGAGCCCCCATGTTCCTGCACTTTGTTGACGTAGCCCAACATGTT
mtDNA_final    AACATGTTCTCTTGTGGCTCGTCCGAGTCGCTCGGCTTCGCCTCGCTCCTCGTCCCGAGCCCCCATGTTCCTGCACTTTGTTGACGTAGCCCAACATGTT

                       110       120       130       140       150       160       170       180       190       200
               ....|....|....|....|....|....|....|....|....|....|....|....|....|....|....|....|....|....|....|
realigned-end  CTCTCAACATGTTCTCTTAACATGTTCTCCTAACATGTTCGCAGAGACAACATGTTCTCTTGTGGCTCGTCCGAGTCGCTCGGCTTCGCCTCGCTCCTCG
mtDNA_final    CTCTCAACATGTTCTCTTAACATGTTCTCCTAACATGTTCGCAGAGACAACATGTTCTCTTGTGGCTCGTCCGAGTCGCTCGGCTTCGCCTCGCTCCTCG

                       210       220       230       240       250       260       270       280       290       300
               ....|....|....|....|....|....|....|....|....|....|....|....|....|....|....|....|....|....|....|
realigned-end  TCCCGAGCCCCCATGTTCCTGCACTTTGTTGAGATGCTTTCCCAAAGCCCCAACATGTTCTCTTAACATGTTCTCTTAACATGTTCTCTTGTGGCTCGTC
mtDNA_final    TCCCGAGCCCCCATGTTCCTGCACTTTGTTGAGATGCTTTCCCAAAGCCCCAACATGTTCTCTTAACATGTTCTCTTAACATGTTCTCTTGTGGCTCGTC

                       310       320       330       340       350       360       370       380       390       400
               ....|....|....|....|....|....|....|....|....|....|....|....|....|....|....|....|....|....|....|
realigned-end  CGAGTCGCTCGGCTTCGCCTCGCTCCTCGTCCCGAGCCCCCATGGCAATGCCTACAGGCCCAGGACTGCCCCTTCGGGGCAGATACTACATGTTCTCTTG
mtDNA_final    CGAGTCGCTCGGCTTCGCCTCGCTCCTCGTCCCGAGCCCCCATGGCAATGCCTACAGGCCCAGGACTGCCCCTTCGGGGCAGATACTACATGTTCTCTTG

                       410       420       430       440       450       460       470       480       490       500
               ....|....|....|....|....|....|....|....|....|....|....|....|....|....|....|....|....|....|....|
realigned-end  TGGCTCGTCCGAGTCGCTCGGCTTCGCCTCGCTCCTCGTACGAGCCAGGTCATGTCCGTAACCCGTTCCCGCGAACCGTTCCCCCGAACCGTTCCCCCGA
mtDNA_final    TGGCTCGTCCGAGTCGCTCGGCTTCGCCTCGCTCCTCGTACGAGCCAGGTCATGTCCGTAACCCGTTCCCGCGAACCGTTCCCCCGAACCGTTCCCCCGA

                       510       520       530       540       550       560       570       580       590       600
               ....|....|....|....|....|....|....|....|....|....|....|....|....|....|....|....|....|....|....|
realigned-end  ACCGTTCCCCCGAACCGTTCCCCCAACCGTTCCCCCGAACCGTTCCCCCGAACCGTTCCCCCGAACCGTTCCCCCGAACCGTTCCCCCGAACCGTTCCCC
mtDNA_final    ACCGTTCCCCCGAACCGTTCCCCCAACCGTTCCCCCGAACCGTTCCCCCGAACCGTTCCCCCGAACCGTTCCCCCGAACCGTTCCCC-GAACCGTTCCCC

                       610       620       630       640       650       660       670       680       690       700
               ....|....|....|....|....|....|....|....|....|....|....|....|....|....|....|....|....|....|....|
realigned-end  CGAACCGTTCCCCC-AACCGTTCCCCCGAACCGTTCCCCCGAACCGTTCCCCCGAACCGTTCCCCCGAACCGTTCCCCCGAACCGTTCCCCCAGTCCAAC
mtDNA_final    --AACCGTTCCCCCGAACCGTTCCCCCGAACCGTTCCCCCGAACCGTTCCCCCGAACCGTTCCCCCGAACCGTTCCCCC-------------AGTCCAAC

                       710       720       730       740       750       760       770       780       790       800
               ....|....|....|....|....|....|....|....|....|....|....|....|....|....|....|....|....|....|....|
realigned-end  CCCCGCCCCGTCAACCGTCCCCCGAACCGTTCCCCCGAACCGTTCCCCCAGGGGGCCATGAGTCCGAGATTGGTTGGCAAGACATGATTCTGCACCCCGC
mtDNA_final    CCCCGCCCCGTCAACCGTCCCCCGAACCGTTCCCCCGAACCGTTCCCCCAGGGGGCCATGAGTCCGAGATTGGTTGGCAAGACATGATTCTGCACCCCGC

                       810       820       830       840       850       860       870       880       890       900
               ....|....|....|....|....|....|....|....|....|....|....|....|....|....|....|....|....|....|....|
realigned-end  TAGTTCCCACATGAGTCTGCTAAGTATTTATGATTCTGCTAAGTCTTGCCCCAACATGTTCTCTTGTGGCTCGTCCGAGTCGCTCGGCTTCGCCTCGCTC
mtDNA_final    TAGTTCCCACATGAGTCTGCTAAGTATTTATGATTCTGCTAAGTCTTGCCCCAACATGTTCTCTTGTGGCTCGTCCGAGTCGCTCGGCTTCGCCTCGCTC

                       910       920       930       940       950       960       970       980       990      1000
               ....|....|....|....|....|....|....|....|....|....|....|....|....|....|....|....|....|....|....|
realigned-end  CTCGTCCTTCGCCTCATGTTCTCTTGTGGCTCGTCCGAGTCGCTCGGCTTCGCCTCGCTCCTCGTCCTTCGCCTCATGTTCTCTTAACATGTTCTCTTGT
mtDNA_final    CTCGTCCTTCGCCTCATGTTCTCTTGTGGCTCGTCCGAGTCGCTCGGCTTCGCCTCGCTCCTCGTCCTTCGCCTCATGTTCTCTTAACATGTTCTCTTGT

                      1010      1020      1030      1040      1050      1060      1070      1080      1090      1100
               ....|....|....|....|....|....|....|....|....|....|....|....|....|....|....|....|....|....|....|
realigned-end  GGCTCGTCCGAGTCGCTCGGCTTCGCCTCGCTCCTCGTCCTTCGCCTCATGTTCTCTTAACATGTTCTCTTGTGGCTCGTCCGAGTCGCTCGGCTTCGCC
mtDNA_final    GGCTCGTCCGAGTCGCTCGGCTTCGCCTCGCTCCTCGTCCTTCGCCTCATGTTCTCTTAACATGTTCTCTTGTGGCTCGTCCGAGTCGCTCGGCTTCGCC

                      1110      1120      1130      1140      1150      1160      1170      1180      1190      1200
               ....|....|....|....|....|....|....|....|....|....|....|....|....|....|....|....|....|....|....|
realigned-end  TCGCTCCTCGTACGAGCCAGGTCATGTCCGTAACCCGTTCCCGCGAACCGTTCCCCCGAACCGTTCCCCCGAACCGTTCCCCCGAACCGTTCCCCCGAAC
mtDNA_final    TCGCTCCTCGTACGAGCCAGGTCATGTCCGTAACCCGTTCCCGCGAACCGTTCCCCCGAACCGTTCCCCCGAACCGTTCC-CCGAACCGTTCCCCCGAAC

                      1210      1220      1230      1240      1250      1260      1270      1280      1290      1300
               ....|....|....|....|....|....|....|....|....|....|....|....|....|....|....|....|....|....|....|
realigned-end  CGTTCCCCCGAACCGTTCCCCCGAACCGTTCCCCCGAACCGTTCCCCCGAACCGTTCCCCCAGTCCAACCCCCGCCCCGTCAACCGT
mtDNA_final    CGTTCCCCCGAACCGTTCCCCCGAACCGTTCCCCCGAACCGTTCCCCCGAACCGTTCC-CCAGTCCAACC-CCGCCCCGTCAACCGT

                      1310      1320      1330      1340      1350      1360      1370      1380      1390      1400
               ....|....|....|....|....|....|....|....|....|....|....|....|....|....|....|....|....|....|....|
realigned-end  CCCCCGAACCGTTCCCCCGAACCGTT-CCCCCAGGGGGCCATGAGTCCGAGATTGGTTGGCAAGACATGATTCTGCACCCCGCTAGTTCCCACATGAGTC
mtDNA_final    CCCCCGAACCGTTC-CCCGAACCGTTTCCCCCAGGGGGCCATGAGTCCGAGATTGGTTGGCAAGACATGATTCTGCACCCCGCTAGTTCCCACATGAGTC

                      1410      1420      1430      1440      1450      1460      1470      1480      1490      1500
               ....|....|....|....|....|....|....|....|....|....|....|....|....|....|....|....|....|....|....|
realigned-end  TGCTAAGTATTTATGATATTCGGATGCTTTCCCAAAGCCCCAACATGTTCTCTTGTGGCTCGTCCGAGTCGCTCGGCTTCGCCTCGCTCCTCGTCCTTCG
mtDNA_final    TGCTAAGTATTTATGATATTCGGATGCTTTCCCAAAGCCCCAACATGTTCTCTTGTGGCTCGTCCGAGTCGCTCGGCTTCGCCTCGCTCCTCGTCCTTCG

                      1510      1520      1530      1540      1550      1560      1570      1580      1590      1600
               ....|....|....|....|....|....|....|....|....|....|....|....|....|....|....|....|....|....|....|
realigned-end  CCTCATGTTCTCTTAACATGTTCTCTTGTGGCTCGTCCGAGTCGCTCGGCTTCGCCTCGCTCCTCGTCCCGAGCCCCCATGTTCCTGCACTTTGTTGACG
mtDNA_final    CCTCATGTTCTCTTAACATGTTCTCTTGTGGCTCGTCCGAGTCGCTCGGCTTCGCCTCGCTCCTCGTCCCGAGCCCCCATGTTCCTGCACTTTGTTGACG

                      1610      1620      1630      1640      1650      1660      1670      1680      1690      1700
               ....|....|....|....|....|....|....|....|....|....|....|....|....|....|....|....|....|....|....|
realigned-end  TAGCCCAACATGTTCTATCAACATGTTCTCTTGTGGCTCGTCCGAGTCGCTCGGCTTCGCCTCGCTCCTCGTCCTTCGCCTCATGTTCTCTTGTGGCTCG
mtDNA_final    TAGCCCAACATGTTCTATCAACATGTTCTCTTGTGGCTCGTCCGAGTCGCTCGGCTTCGCCTCGCTCCTCGTCCTTCGCCTCATGTTCTCTTGTGGCTCG

                      1710      1720      1730      1740      1750      1760      1770      1780      1790      1800
               ....|....|....|....|....|....|....|....|....|....|....|....|....|....|....|....|....|....|....|
realigned-end  TCCGAGTCGCTCGGCTTCGCCTCGCTCCTCGTCCTTCGCCTCATGTTCTCTTAACATGTTCTCTTGTGGCTCGTCCGAGTCGCTCGGCTTCG-----CTC
mtDNA_final    TCCGAGTCGCTCGGCTTCGCCTCGCTCCTCGTCCTTCGCCTCATGTTCTCTTAACATGTTCTCTTGTGGCTCGTCCGAGTCGCTCGGCTTCGCCTCGCTC

                      1810      1820      1830      1840      1850      1860      1870      1880      1890      1900
               ....|....|....|....|....|....|....|....|....|....|....|....|....|....|....|....|....|....|....|
realigned-end  CTCGTCCCGAGCCCCCATGTTCCTGCACATGTTCTCTTGTGGCTCGTCCGAGTCGCTCGGCTTCGCCTCGCTCCTCGTCCTTCGCCTCATGTTCTCTTAA
mtDNA_final    CTCGTCCCGAGCCCCCATGTTCCTGCACATGTTCTCTTGTGGCTCGTCCGAGTCGCTCGGCTTCGCCTCGCTCCTCGTCCTTCGCCTCATGTTCTCTTAA

                      1910      1920      1930      1940      1950      1960      1970      1980      1990      2000
               ....|....|....|....|....|....|....|....|....|....|....|....|....|....|....|....|....|....|....|
realigned-end  CATGTTCTCTTGTGGCTCGTCCGAGTCGCTCGGCTTCGCCTCGCTCCTCGTCCTTCGCCTCATGTTCTCTTAACATGTTCTCTTGTGGCTCGTCCGAGTC
mtDNA_final    CATGTTCTCTTGTGGCTCGTCCGAGTCGCTCGGCTTCGCCTCGCTCCTCGTCCTTCGCCTCATGTTCTCTTAACATGTTCTCTTGTGGCTCGTCCGAGTC

                      2010      2020      2030      2040      2050      2060      2070      2080      2090      2100
               ....|....|....|....|....|....|....|....|....|....|....|....|....|....|....|....|....|....|....|
```

```
realigned-end   GCTCGGCTTCGCCTCGCTCCTCGTCCTTCGCCTCATGTTCTCTTAACATGTTCTCTTGTGGCTCGTCCGAGTCGCTCGGCTTCGCCTCGCTCCTCGTCCC
mtDNA_final     GCTCGGCTTCGCCTCGCTCCTCGTCCTTCGCCTCATGTTCTCTTAACATGTTCTCTTGTGGCTCGTCCGAGTCGCTCGGCTTCGCCTCGCTCCTCGTCCC

                      2110      2120      2130      2140      2150      2160      2170      2180      2190      2200
                ....|....|....|....|....|....|....|....|....|....|....|....|....|....|....|....|....|....|....|....|
realigned-end   GAGCCCCCATGGCAATGCCTACAGGCCCAGGACTGCCCCTTCGGGGCAGATACNACATGTTCTCTTGTGGCTCGTCCGAGTCGCTCGGCTTCGCCTCGCT
mtDNA_final     GAGCCCCCATGGCAATGCCTACAGGCCCAGGACTGCCCCTTCGGGGCAGATACTACATGTTCTCTTGTGGCTCGTCCGAGTCGCTCGGCTTCGCCTCGCT

                      2210      2220      2230      2240      2250      2260      2270      2280      2290      2300
                ....|....|....|....|....|....|....|....|....|....|....|....|....|....|....|....|....|....|....|....|
realigned-end   CCTCGTACGAGCCAGGTCATGTCCGTAACCCGTTCCCGCGAACCGTTCCCCCGAACCGTTCCCCCGAACCGTTCCCCCAGTCCAACC-CCGCCCCGTCAA
mtDNA_final     CCTCGTACGAGCCAGGTCATGTCCGTAACCCGTTCCCGCGAACCGTTCCCCCGAACCGTTCCCCCGAACCGTTCCCCCAGTCCAACCCCCGCCCCGTCAA

                      2310      2320      2330      2340      2350      2360      2370      2380      2390      2400
                ....|....|....|....|....|....|....|....|....|....|....|....|....|....|....|....|....|....|....|....|
realigned-end   CCGTCCC-CGAACCGTTCCC-CGAACCGTTCCC-C-GGGGGCCATGAGTCCGAGATTGGTTGGCAAGACATGATTCTGCACCCCGCTAGTTCCCACATGA
mtDNA_final     CCGTCCCCCGAACCGTTCCCCCGAACCGTTCCCCCAGGGGGCCATGAGTCCGAGATTGGTTGGCAAGACATGATTCTGCACCCCGCTAGTTCCCACATGA

                      2410      2420      2430      2440      2450      2460      2470      2480      2490      2500
                ....|....|....|....|....|....|....|....|....|....|....|....|....|....|....|....|....|....|....|....|
realigned-end   TTCTGCACCCCGCTAGTTCCCACATGATTCTGCAAAGTATATCCAAGCGCTGATCTGGGTGTTCTCTTAACATGTTCTCTTGTGGCTCGTCCGAGTCGCT
mtDNA_final     TTCTGCACCCCGCTAGTTCCCACATGATTCTGCAAAGTATATCCAAGCGCTGATCTGGGTGTTCTCTTAACATGTTCTCTTGTGGCTCGTCCGAGTCGCT

                      2510      2520      2530      2540      2550      2560      2570      2580      2590      2600
                ....|....|....|....|....|....|....|....|....|....|....|....|....|....|....|....|....|....|....|....|
realigned-end   CGGCTTCGCCTCGCTCCTCGTCTTCGCCTAGCTCCTCGTACCTGATCCAAGCGCTGATCTTCCAAGCGCTGATCTGGGCCATCCGGGAGCTTGGGCTTGT
mtDNA_final     CGGCTTCGCCTCGCTCCTCGTCTTCGCCTAGCTCCTCGTACCTGATCCAAGCGCTGATCTTCCAAGCGCTGATCTGGGCCATCCGGGAGCTTGGGCTTGT

                      2610      2620      2630      2640      2650      2660      2670      2680      2690      2700
                ....|....|....|....|....|....|....|....|....|....|....|....|....|....|....|....|....|....|....|....|
realigned-end   GCCCCCCCGTTACCCGTTCCCGTAACCCGTTCCCGTAACCCGTTCCCGCGAACCGTTCCCCCGAACCGTTCCCCCGAACCGTTCCCCCGAACCGTTCCCC
mtDNA_final     GCCCCCCCGTTACCCGTTCCCGTAACCCGTTCCCGTAACCCGTTCCCGCGAACCGTTCCCCCGAACCGTTCCCCCGAACCGTTCCCCCGAACCGTTCCCC

                      2710      2720      2730      2740      2750      2760      2770      2780      2790      2800
                ....|....|....|....|....|....|....|....|....|....|....|....|....|....|....|....|....|....|....|....|
realigned-end   TGAACCGTTCCCCCGAACCGTTCCCCCGAACCGTTCCCCCAACCGTTCCCCCGAACCGTTCCCTCGAACCGTTCCCCCGAACCGTTCCCCCAACCGTTCC
mtDNA_final     TGAACCGTTCCCCCGAACCGTTCCCCCGAACCGTTCCCCCAACCGTTCCCCCGAACCGTTCCCTCGAACCGTT-CCCCGAACCGTTCCCCCAACCGTTCC

                      2810      2820      2830      2840      2850      2860      2870      2880      2890      2900
                ....|....|....|....|....|....|....|....|....|....|....|....|....|....|....|....|....|....|....|....|
realigned-end   CCCGAACCGTTCCCTCAGCAGATACAACATGTTCTCTTGTGGCTCGTCCGAGTCGCTCGGCTTCGCCTCGCTCCTCGTCCTTCGCCTCGCAAGAGGGCAA
mtDNA_final     CCCGAACCGTTCCCTCAGCAGATACAACATGTTCTCTTGTGGCTCGTCCGAGTCGCTCGGCTTCGCCTCGCTCCTCGTCCTTCGCCTCGCAAGAGGGCAA

                      2910      2920      2930      2940      2950      2960      2970      2980      2990      3000
                ....|....|....|....|....|....|....|....|....|....|....|....|....|....|....|....|....|....|....|....|
realigned-end   CGGTAAGAGGGCAACGGTAAGAGGGCAACGGTAAGAGGGCAACGGTAAGAGGGCAACGGTAAGAGGGCAACGGTAAGAGGGCAACGGTAAGAGGGCAACG
mtDNA_final     CGGTAAGAGGGCAACGGTAAGAGGGCAACGGTAAGAGGGCAACGGTAAGAGGGCAACGGTAAGAGGGCAACGGTAAGAGGGCAACGGT--GAGGGCAANN

                      3010      3020      3030      3040      3050      3060      3070      3080      3090      3100
                ....|....|....|....|....|....|....|....|....|....|....|....|....|....|....|....|....|....|....|....|
realigned-end   GTAAGAGGGCAACGGTAAGAGGGCAACGGTAAGAGGGCAACGGTAAGAGGGCAAGCAGAGACAACATGTTCTCTTGTGGCTCGTCCGAGTCGCTCGGCTT
mtDNA_final     NNNNNNNNNCAACGGTAAG-GGGCAACGGTAAGAGGGCAACGGTAAGAGGGCAAGCAGAGACAACATGTTCTCTTGTGGCTCGTCCGAGTCGCTCGGCTT

                      3110      3120      3130      3140      3150      3160      3170      3180      3190      3200
                ....|....|....|....|....|....|....|....|....|....|....|....|....|....|....|....|....|....|....|....|
realigned-end   CGCCTCGCTCCTCGTCCTTCGCCTCATGTTCTCTTAACATGTTCTCTTGTGGCTCGTCCGAGTCGCTCGGCTTCGCCTCGCAAGAGGGCAACGGTAAGAG
mtDNA_final     CGCCTCGCTCCTCGTCCTTCGCCTCATGTTCTCTTAACATGTTCTCTTGTGGCTCGTCCGAGTCGCTCGGCTTCGCCTCGCAAGAGGGCAACGGTAAGAG

                      3210      3220      3230      3240      3250      3260      3270      3280      3290      3300
                ....|....|....|....|....|....|....|....|....|....|....|....|....|....|....|....|....|....|....|....|
realigned-end   GGCAACGGTAAGAGGGCAACGGTAAGAGGGCAACGGTAAGAGGGCAACGGTAAGAGGGCAACGGTAAGAGGGCAACGGTAAGAGGGCAAGCAGAGACAAC
mtDNA_final     GGCAACGGTAAAGGGGCAACGGTAAGAGGGCAACGGTAAGAGGGCAACGGTAA-AGGGCAACGGTAAG-GGGCAACGGTAA-AGGGCAAGCAGAGACAAC

                      3310      3320      3330      3340      3350      3360      3370      3380      3390      3400
                ....|....|....|....|....|....|....|....|....|....|....|....|....|....|....|....|....|....|....|....|
realigned-end   ATGTTCTCTTGTGGCTCGTCCGAGTCGCTCGGCTTCGCCTCGCTCCTCGTCTTCGCCTAGCTCCTCGTACCTGATCCAAGCGCTGATCTTCCAAGCGCTG
mtDNA_final     ATGTTCTCTTGTGGCTCGTCCGAGTCGCTCGGCTTCGCCTCGCTCCTCGTCTTCGCCTAGCTCCTCGTACCTGATCCAAGCGCTGATCTTCCAAGCGCTG

                      3410      3420      3430      3440      3450      3460      3470      3480      3490      3500
                ....|....|....|....|....|....|....|....|....|....|....|....|....|....|....|....|....|....|....|....|
realigned-end   ATCTGGGCCATCCGGGAGCTTGGGCTTGTGCCCCCCCGTTACCCGTTCCCGTAACCCGTTCCCGTAACCCGTTCCCGCGAACCGTTCCCCCGAACCGTTC
mtDNA_final     ATCTGGGCCATCCGGGAGCTTGGGCTTGTGCCCCCCCGTTACCCGTTCCCGTAACCCGTTCCCGTAACCCGTTCCCGCGAACCGTTCCCCCGAACCGTTC

                      3510      3520      3530      3540      3550      3560      3570      3580      3590      3600
                ....|....|....|....|....|....|....|....|....|....|....|....|....|....|....|....|....|....|....|....|
realigned-end   CCCCGAACCGTTCCCCCGAACCGTTCCCCTGAACCGTTCCCCCGAACCGTTCCCCCGAACCGTTCCCCCAACCGTTCCCCCGAACCGTTCCCTCGAACCG
mtDNA_final     CCCCGAACCGTTCCCCCGAACCGTTCCCCTGAACCGTTCCCCCGAACCGTTCCCCCGAACCGTTCCCCCAACCGTTCCCCCGAACCGTTCCCTCGAACCG

                      3610      3620      3630      3640      3650      3660      3670      3680      3690      3700
                ....|....|....|....|....|....|....|....|....|....|....|....|....|....|....|....|....|....|....|....|
realigned-end   TTCCCCCGAACCGTTCCCCCAACCGTTCCCCCGAACCGTTCCCCCGAACCGTTCCCCCGAACCGTCCCCCCGAACCGTTCCCCCGAACCGTTCCCCCGAA
mtDNA_final     TTCCCCCGAACCGTTCCCCCAACCGTTCCCCCGAACCGTTCCCCCGAACCGTTCCCCCGAACCGTCCCCCCGAACCGTTCCCCCGAACCGTTCCCCCGAA

                      3710      3720      3730      3740      3750      3760      3770      3780      3790      3800
                ....|....|....|....|....|....|....|....|....|....|....|....|....|....|....|....|....|....|....|....|
realigned-end   CCGTTCCCCCGAACCGTTCCCCCGAACCGTCCCCCCGAACCGTTCCCCCGAACCGTTCCCCCAGTCCAACCCCCGCCTGATGATACTATGATTCCCATAT
mtDNA_final     CCGTTCCCCCGAACCGTTCCCCCGAACCGTCCCCCCGAACCGTTCCCCCGAACCGTTCCCCCAGTCCAACCCCCGCCTGATGATACTATGATTCCCATAT

                      3810      3820      3830      3840      3850      3860      3870      3880      3890      3900
                ....|....|....|....|....|....|....|....|....|....|....|....|....|....|....|....|....|....|....|....|
realigned-end   CAAATCCTAGTTCCCATCTGATTCTGCTAAGTCTTTCTGATTCTGCTAAGTCTTTCAGATTCTGCTAAGTCTTTCTGATTCTGCACCCCGCTAGTTCCCA
mtDNA_final     CAAATCCTAGTTCCCATCTGATTCTGCTAAGTCTTTCTGATTCTGCTAAGTCTTTCAGATTCTGCTAAGTCTTTCTGATTCTGCACCCCGCTAGTTCCCA

                      3910      3920      3930      3940      3950      3960      3970      3980      3990      4000
                ....|....|....|....|....|....|....|....|....|....|....|....|....|....|....|....|....|....|....|....|
realigned-end   CATGATTCTGCTAAGTATTTATGATTCAGCTAAGTCTTGCCCCAACATGTTCTCTTGTGGCTCGTCCGAGTCGCTCGGCTTCGCCTCGCTCCTCGTCTTC
mtDNA_final     CATGATTCTGCTAAGTATTTATGATTCAGCTAAGTCTTGCCCCAACATGTTCTCTTGTGGCTCGTCCGAGTCGCTCGGCTTCGCCTCGCTCCTCGTCTTC

                      4010      4020      4030      4040      4050      4060      4070      4080      4090      4100
                ....|....|....|....|....|....|....|....|....|....|....|....|....|....|....|....|....|....|....|....|
realigned-end   GCCTAGCTCCTCGTACCTGATCCAAGCGCTGATCTGGGCCATCCGGGAGCTTGGGCTTGTGCCCCACCTGATCCAAGCGCTGATCTGGGCCATCCGGGAG
mtDNA_final     GCCTAGCTCCTCGTACCTGATCCAAGCGCTGATCTGGGCCATCCGGGAGCTTGGGCTTGTGCCCCACCTGATCCAAGCGCTGATCTGGGCCATCCGGGAG
```

```
                      4110      4120      4130      4140      4150      4160      4170      4180      4190      4200
                ....|....|....|....|....|....|....|....|....|....|....|....|....|....|....|....|....|....|....|....|
realigned-end   CTTGGGCTTGTGCCCCC--GTTACCCGTTCCCGTAACCCGTTCCCGTAACCCGTTCCCGCGAACCGTTCCCCCGAACCGTTCCCCCGAACCGTTCCCCCG
mtDNA_final     CTTGGGCTTGTGCCCCCCCGTTACCCGTTCCCGTAACCCGTTCCCGTAACCCGTTCCCGCGAACCGTTCCCCCGAACCGTTCCCCCGAACCGTTCCCCCG

                      4210      4220      4230      4240      4250      4260      4270      4280      4290      4300
                ....|....|....|....|....|....|....|....|....|....|....|....|....|....|....|....|....|....|....|
realigned-end   AACCGTTCCCCTGAACCGTTCCCCCGAACCGTTCCCCCGAACCGTTCCCCCAACCGTTCCCCCGAACCGTTCCCTCGAACCGTTCCCCCGAACCGTTCCC
mtDNA_final     AACCGTTCCCCTGAACCGTTCCCCCGAACCGTTCCCCCGAACCGTTCCCCCAACCGTTCCCCCGAACCGTTCCCTCGAACCGTTCCCCCGAACCGTTCCC

                      4310      4320      4330      4340      4350      4360      4370      4380      4390      4400
                ....|....|....|....|....|....|....|....|....|....|....|....|....|....|....|....|....|....|....|
realigned-end   CCAACCGTTCCCCCGAACCGTTCCCCCGAACCGTTCCCCCAGTCCAACCCCCGCCCCGTCAACCGTCCCCCGAACCGTTCCCCCGAACCGTTCCCCCAGG
mtDNA_final     CCAACCGTTCCCCCGAACCGTTCCCCCGAACCGTTCCCCCAGTCCAACCCCCGCCCCGTCAACCGTCCCCCGAACCGTTCCCCCGAACCGTTCCCCCAGG

                      4410      4420      4430      4440      4450      4460      4470      4480      4490      4500
                ....|....|....|....|....|....|....|....|....|....|....|....|....|....|....|....|....|....|....|
realigned-end   GGGCCATGAGTCCGAGATTGGTTGGCAAGACATGATTCTGCACCCCGCTAGTTCCCACATGATTCTGCACCCCGCTAGTTCCCACATGATTCTGCACCCC
mtDNA_final     GGGCCATGAGTCCGAGATTGGTTGGCAAGACATGATTCTGCACCCCGCTAGTTCCCACATGATTCTGCACCCCGCTAGTTCCCACATGATTCTGCACCCC

                      4510      4520      4530      4540      4550      4560      4570      4580      4590      4600
                ....|....|....|....|....|....|....|....|....|....|....|....|....|....|....|....|....|....|....|
realigned-end   GCTAGTTCCCA--TGATTCTGCAAAGTATTGTTACGCTAACTGTTCCTGAGCTGCCCGGTAGCTCCCACGTAATTCCGCTAAGTATTAATGATTATGCAC
mtDNA_final     GCTAGTTCCCACATGATTCTGCAAAGTATTGTTACGCTAACTGTTCCTGAGCTGCCCGGTAGCTCCCACGTAATTCCGCTAAGTATTAATGATTATGCAC

                      4610      4620      4630      4640      4650      4660      4670      4680      4690      4700
                ....|....|....|....|....|....|....|....|....|....|....|....|....|....|....|....|....|....|....|
realigned-end   CCCGCTAGTTCCCACATGAGTCTGCTAAGTATTTATGAGTCTGCTAAGTCTTGCCCCAACATGTTCTCTTGTGGCTCGTCCGAGTCGCTCGGCTTCGCCT
mtDNA_final     CCCGCTAGTTCCCACATGAGTCTGCTAAGTATTTATGAGTCTGCTAAGTCTTGCCCCAACATGTTCTCTTGTGGCTCGTCCGAGTCGCTCGGCTTCGCCT

                      4710      4720      4730      4740      4750      4760      4770      4780      4790      4800
                ....|....|....|....|....|....|....|....|....|....|....|....|....|....|....|....|....|....|....|
realigned-end   CGCTCCTCGTCTTCGCCTAGCTCCTCGTGATACAACATGTTCTCTTGTGGCTCGTCCGAGTCGCTCGGCTTCGCCTCGCTCCTCGTCCCGAGCCCCCATG
mtDNA_final     CGCTCCTCGTCTTCGCCTAGCTCCTCGTGATACAACATGTTCTCTTGTGGCTCGTCCGAGTCGCTCGGCTTCGCCTCGCTCCTCGTCCCGAGCCCCCATG

                      4810      4820      4830      4840      4850      4860      4870      4880      4890      4900
                ....|....|....|....|....|....|....|....|....|....|....|....|....|....|....|....|....|....|....|
realigned-end   TTCCTGCACTTTGTTGACGTAGCCCAACATGTTCTATCAACATGTTCTCTTGTGGCTCGTCCGAGTCGCTCGGCTTCGCCTCGCTCCTCGTCCTTCGCCT
mtDNA_final     TTCCTGCACTTTGTTGACGTAGCCCAACATGTTCTATCAACATGTTCTCTTGTGGCTCGTCCGAGTCGCTCGGCTTCGCCTCGCTCCTCGTCCTTCGCCT

                      4910      4920      4930      4940      4950      4960      4970      4980      4990      5000
                ....|....|....|....|....|....|....|....|....|....|....|....|....|....|....|....|....|....|....|
realigned-end   CATGTTCTCTTGTGGCTCGTCCGAGTCGCTCGGCTTCGCCTCGCTCCTCGTCCTTCGCCTCATGTTCTCTTAACATGTTCTCTTGTGGCTCGTCCGAGTC
mtDNA_final     CATGTTCTCTTGTGGCTCGTCCGAGTCGCTCGGCTTCGCCTCGCTCCTCGTCCTTCGCCTCATGTTCTCTTAACATGTTCTCTTGTGGCTCGTCCGAGTC

                      5010      5020      5030      5040      5050      5060      5070      5080      5090      5100
                ....|....|....|....|....|....|....|....|....|....|....|....|....|....|....|....|....|....|....|
realigned-end   GCTCGGCTTCGCCTCGCTCCTCGTCCCGAGCCCCCATGGCAATGCCTACAGGCCCAGGACTGCCCCTTCGGGGCAGAGACAACATGTTCTCTTGTGGCTC
mtDNA_final     GCTCGGCTTCGCCTCGCTCCTCGTCCCGAGCCCCCATGGCAATGCCTACAGGCCCAGGACTGCCCCTTCGGGGCAGAGACAACATGTTCTCTTGTGGCTC

                      5110      5120      5130      5140      5150      5160      5170      5180      5190      5200
                ....|....|....|....|....|....|....|....|....|....|....|....|....|....|....|....|....|....|....|
realigned-end   GTCCGAGTCGCTCGGCTTCGCCTCGCTCCTCGTCTTCGCCTAGCTCCTCGTACCTGATCCAAGCGCTGATCTGGGCCATCCGGGAGCTTGGGCTTGTGCC
mtDNA_final     GTCCGAGTCGCTCGGCTTCGCCTCGCTCCTCGTCTTCGCCTAGCTCCTCGTACCTGATCCAAGCGCTGATCTGGGCCATCCGGGAGCTTGGGCTTGTGCC

                      5210      5220      5230      5240      5250      5260      5270      5280      5290      5300
                ....|....|....|....|....|....|....|....|....|....|....|....|....|....|....|....|....|....|....|
realigned-end   CCACCTGATCCAAGCGCTGATCTGGGCCATCCGGGAGCTTGGGCTTGTGCCCCCCCGTTACCCGTTCCCGTAACCCGTTCCCGTAACCCGTTCCCGCGAA
mtDNA_final     CCACCTGATCCAAGCGCTGATCTGGGCCATCCGGGAGCTTGGGCTTGTGCCCCCCCGTTACCCGTTCCCGTAACCCGTTCCCGTAACCCGTTCCCGCGAA

                      5310      5320      5330      5340      5350      5360      5370      5380      5390      5400
                ....|....|....|....|....|....|....|....|....|....|....|....|....|....|....|....|....|....|....|
realigned-end   CCGTTCCCCCGAACCGTTCCCCCGAACCGTTCCCCCGAACCGTTCC-CTGAACCGTTCCC-CGAACCGTTCCCCGAACCGTTCCCCCAACCGTTCCCCGA
mtDNA_final     CCGTTCCCCCGAACCGTTCCCCCGAACCGTTCCCCCGAACCGTTCCCCTGAACCGTTCCCCCGAACCGTTCCCCGAACCGTTCC-CCAACCGTTCCCCGA

                      5410      5420      5430      5440      5450      5460      5470      5480      5490      5500
                ....|....|....|....|....|....|....|....|....|....|....|....|....|....|....|....|....|....|....|
realigned-end   ACCGTTCCCTCGAACCGTTCCCCGAACCGTTCCCCCAACCGTTCCCCCGAACCGTTCCCTCAGCAGATACAACATGTTCTCTTGTGGCTCGTCCGAGTCG
mtDNA_final     ACCGTTC-CTCGAACCGTTCCCCGAACCGTTCC-CCAACCGTTCCCCCGAACCGTTCCCTCAGCAGATACAACATGTTCTCTTGTGGCTCGTCCGAGTCG

                      5510      5520      5530      5540      5550      5560      5570      5580      5590      5600
                ....|....|....|....|....|....|....|....|....|....|....|....|....|....|....|....|....|....|....|
realigned-end   CTCGGCTTCGCCTCGCTCCTCGTCCTTCGCCTCGCAAGAGG-CAACGGTAAGAG-ACAACGGTAAGAGGGCAACGGTAAGAGGGCAACGGTAAGAGGGCA
mtDNA_final     CTCGGCTTCGCCTCGCTCCTCGTCCTTCGCCTCGCAAGAGGGCAACGGTAAGAGGGCAACGGTAAGAGGGCAACGGTAAGAGGGCAACGGTAAGAGGGCA

                      5610      5620      5630      5640      5650      5660      5670      5680      5690      5700
                ....|....|....|....|....|....|....|....|....|....|....|....|....|....|....|....|....|....|....|
realigned-end   ACGGTAAGAGGGCAACGGTAAGAGGGCAA----------------------------------------------------------------------G
mtDNA_final     ACGGTAAGAGGGCAACGGTAAGAGGGCAACGGTAAGAGGGCAACGGTAAGAGGGCAACGGTAAGAGGGCAACGGTAAGAGGGCAACGGTAAGAGGGCAAG

                      5710      5720      5730      5740      5750      5760      5770      5780      5790      5800
                ....|....|....|....|....|....|....|....|....|....|....|....|....|....|....|....|....|....|....|
realigned-end   CAGAGACAACATGTTCTCTTGTGGCTCGTCCGAGTCGCTCGGCTTCGCCTCGCTCCTCGTCCTTCGCCTCA
mtDNA_final     CAGAGACAACATGTTCTCTTGTGGCTCGTCCGAGTCGCTCGGCTTCGCCTCGCTCCTCGTCCTTCGCCTCATGTTCTCTTGTGGCTCGTCCGAGTCGCTC
```

**Fig. S8** Alignment of the highly similar terminal regions of the originally assembled linear mitogenome contig. The sequence similarity of the termini is 97.7 % (along the alignment containing 5,771 nucleotide positions). The sequence marked here as "mtDNA_final" represents the variant of the terminus sequence that was retained in the final putatively circular-mapping full mitogenome sequence.

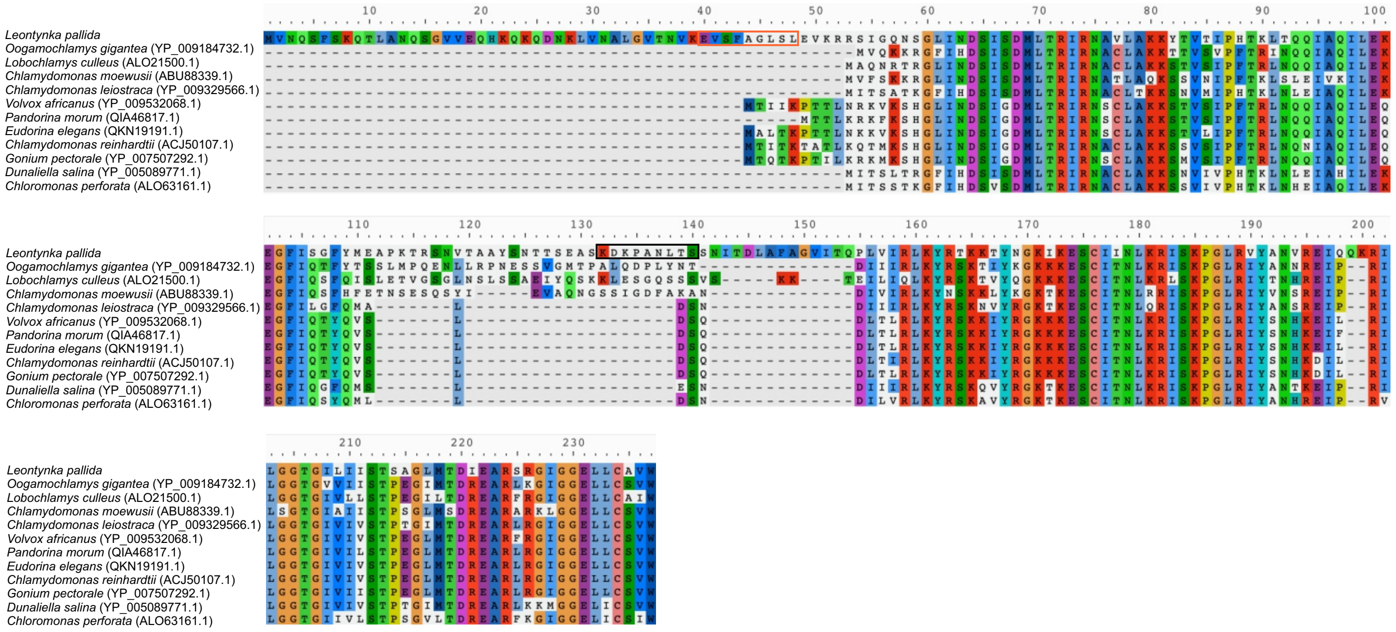


**Fig. S9** Occurrence of the "variant 8" repeat (translated in reading frame +0 as KDKPANLTS and -0 as KEVSFAGLSL; both boxed in colour) in a variable region of protein sequence of the ribosomal protein Rps8 from *Leontynka pallida* (full protein alignment together with representatives of other chlamydomonadalean algae).
