## Additional file 2: Supplementary Notes and supplementary Methods for "A new lineage of non-photosynthetic green algae with extreme organellar genomes"

**This file includes supplementary notes, supplementary methods, and relevant supplementary references.**

### Supplementary notes

**Note S1** Taxonomic descriptions

Here we follow the rules and recommendations of The International Code of Nomenclature (ICN) for algae, fungi, and plants [13].

Chlamydomonadales *incertae sedis*

***Leontynka* Pánek, Barcytė, Zadrobílková, Čepička, Eliáš, gen. nov.**

DESCRIPTION: Cells colourless, unicellular, biflagellate, ovoid to almost spherical, or ellipsoidal to ellipsoidal-cylindrical. Cell wall smooth, robust, and with papilla. Two apical contractile vacuoles present. Without photosynthetically active chloroplast; instead, highly distorted but essentially cup-shaped leucoplasts with large starch grains present. Pyrenoid absent. One to two eyespots. Nucleus central or slightly posterior. Lipid droplets present. Asexual reproduction through two or four zoospores. Sexual reproduction not observed.

The genus is morphologically similar to *Polytoma* Ehrenberg, but phylogenetic analyses based on the 18S rRNA gene do not show a close relationship of *Leontynka* spp. to any *Polytoma* spp. (including the type species, *Polytoma uvella*). See also the comprehensive differential diagnoses in Note S4.

ETYMOLOGY: In honour to "Leontýnka", the ghost of a little girl dressed in white, one of the main characters of the Czechoslovakian children's film "Long Live Ghosts!" (1977); inspired by *Rusalka*, a different chlamydomonadalean alga named after a similar esoteric fictional character from Czech mythology. Considered of feminine gender.

TYPE SPECIES: *Leontynka pallida* sp. nov.

***Leontynka pallida*** **Pánek, Barcytė, Zadrobílková, Čepička, Eliáš, sp. nov.**
DESCRIPTION: Cells 12.0–20.0 µm long and 8.0–17.0 µm wide, ovoid to broadly rounded, or almost spherical. Posterior end always broader than the anterior. Cell wall robust and smooth, with a low but prominent keel-shaped papilla, with two isokont flagella approximately as long as the cell. Two apical contractile vacuoles present. Leucoplasts cup-shaped, consisting of separate distorted compartments, each containing a single huge or two smaller starch grains. Lipid droplets present. Eyespot large, ellipsoidal or roundish, usually in the lateral middle or anterior end of the leucoplast, in the last or second third of the anterior cell end, often hidden. A second eyespot with varying position may be present. Nucleus central or slightly posterior with a prominent nucleolus. Asexual reproduction through two or four zoospores. Sexual reproduction not observed.

HOLOTYPE: A mounted specimen (resin block) derived from the strain AMAZONIE has been deposited in the National Museum, Prague, Czech Republic; inventory number P6E 5164.

TYPE LOCALITY: Explorer´s Inn, Tambopata province, Amazon rainforest, Peru, South America.

HABITAT: Hypoxic or microoxic freshwater sediments.

ETYMOLOGY: L. fem. adj. *pallida* – pale.

NOTES: Figures 2a-e show the morphology of the authentic strain. DNA sequences are available in GenBank: OM501587.1 (partial 18S rDNA + ITS1 + 5.8S rDNA + ITS2 region); OM479425.1 (complete plastid genome); OM479424.1 (complete mitochondrial genome).

***Leontynka elongata*** **Pánek, Barcytė, Zadrobílková, Čepička, Eliáš sp. nov.**
DESCRIPTION: Cells 12.0–20.0 µm long and 6.0–14.0 µm wide, ellipsoidal, ellipsoidal-cylindrical or ovoid. Posterior end may be empty. Cell wall thin and smooth, with a prominent keel-shaped papilla, with two isokont flagella approximately as long as the cell. Two apical contractile vacuoles present. Carved cup-shaped leucoplast composed of separate compartments containing one to two starch grains. Lipid droplets present. Eyespot large, ellipsoidal or roundish, usually in the lateral anterior end of the leucoplast, in the first or second third of the anterior cell end, often hidden. A second eyespot can be present. Nucleus central or slightly posterior. Mitochondria contain discoidal cristae. Asexual reproduction through two or four zoospores. Sexual reproduction not observed.

Differs from the closely related *Leontynka pallida* in morphology (cell shape, position of the eyespot), ultrastructure (different mitochondrial cristae), and in 18S and ITS2 rDNA sequences, including compensatory base changes in helix II of the ITS2 structure.

HOLOTYPE: A mounted specimen (resin block) derived from the strain MBURUCU has been deposited in the National Museum, Prague, Czech Republic; inventory number P6E 5165.

TYPE LOCALITY: Mburucuyá, Argentina, South America.

HABITAT: Hypoxic or microoxic freshwater sediments.

ETYMOLOGY: From L. fem. perf. pass. part. *elongata* – elongated (here used as an adjective).

NOTES: Figures 2f-j show the morphology of the authentic strain. DNA sequences are available in GenBank: OM501588.1 (partial 18S rDNA + ITS1 + 5.8S rDNA + ITS2 region).

**Note S2** Further details on the morphology and ultrastructure of *Leontynka* spp.

The cultured cells of *L. pallida* were ovoid and often spherical to subspherical, 11.5-19.7 µm long and 8.0-17.3 µm wide, with an average length of 15.3 (± 1.8) µm and an average width of 11.5 (± 2.0) µm (figures 2a-c and S4). The two flagella were approximately of the same length as the cell (figure S4c). The cells contained a prominent ellipsoidal or slightly roundish eyespot of variable position but usually located in the lateral middle region of the plastid (figures 2a and S4a). Sometimes the eyespot was also present in the lateral anterior end of the plastid, in the second or the last third of the anterior cell end (figure S4b, e). Occasionally, two eyespots were present within a single cell (figure 2b).

In contrast, *L. elongata* had ellipsoidal or broadly ellipsoidal to almost cylindrical, or ovoid cells with rounded ends. Cells were 12.5-19.5 µm long and 6.0-14.0 µm wide, with an average length of 15.6 (± 1.6) µm and an average width of 8.9 (± 1.5) µm (figures 2f-h and S5). The cell posterior was always slightly broader than the anterior. Both flagella were approximately as long as the cell. An ellipsoidal or roundish eyespot was present in the first or second third of the anterior cell end (figures 2f-h and S5a, c-f, h). As in *L. pallida*, a second eyespot was also occasionally present that exhibited a variable position (figure S5d, e).

Cells of both species were covered by a continuous cell wall consisting of two electron-dense layers. An amorphous material was present in the space between the cell wall and the plasma membrane (figures 2j and S6e). The cell wall was dissociated from the protoplast (figures 2d, e, i, j; figure S6) and numerous small electron-dense globules were sometimes scattered in between (figures 2d, e, and S6f). The TEM confirmed the presence of the prominent keel-shaped papilla in both species (figures S6d and g), and a directly opposed flagellar apparatus orientation (figure S6d). Leucoplasts were highly convoluted (figures S6a and h), and were bounded by two membranes (figure S6c). When present, cytoplasmic lipid droplets were scattered among plastidial "bridges" (figures S6b and h). Electron-dense particles were also commonly present within the protoplast (figures 2d, I, and S6a-c, g, h).

**Note S3** Further details on various kinds of repeats in the plastome of *L. pallida*

The imperfect palindrome “CAAACCAGT|AA|ACTGGTTAG” together with the interspersed repeat “TAACTAAACTTC” constitute an abundant tandem composite repeat that is present in several clusters across the plastome. As an example, one such cluster (2,132 bp) is annotated below (positions 26,098-28,230 of the arbitrarily linearized *L. pallidum* plastid genome sequence, GenBank accession number OM479425.1). The regions in yellow correspond to individual copies of the imperfect palindrome, the regions in black represent copies of the interspersed repeat.

“CAAACCAGTAAACTGGTTAGTAACTAAACTTCCAAACCAGTAAACTGGTTAGTAACTAAACTTCCAAAC
CAGTAAACTGGTTAGTAACTAAACTTCCAAACCAGTAAACTGGTTAGTAACTAAACTTCCAAACCAGTAA
ACTGGTTAGTAACTAAACTTCCAAACCAGTAAACTGGTTAGTAACTAAACTTCCAAACCAGTAAACTGGT
TAGTAACTAAACTTCCAAACCAGTAAACTGGTTAGTAACTAAACTTCCAAACCAGTAAACTGGTTAGTAA
CTAAACTTCCAAACCAGTAAACTGGTTAGTAACTAAACTTCCAAACCAGTAAACTGGTTAGTAACTAAAC
TTCCAAACCAGTAAACTGGTTAGTAACTAAACTTCCAAACCAGTAAACTGGTTAGTAACTAAACTTCCAA
ACCAGTAAACTGGTTAGTAACTAAACTTCCAAACCAGTAAACTGGTTAGTAACTAAACTTCCAAACCAGT
AAACTGGTTAGTAACTAAACTTCCAAACCAGTAAACTGGTTAGTAACTAAACTTCCAAACCAGTAAACTG
GTTAGTAACTAAACTTCCAAACCAGTAAACTGGTTAGTAACTAAACTTCCAAACCAGTAAACTGGTTAGT
AACTAAACTTCCAAACCAGTAAACTGGTTAGTAACTAAACTTCCAAACCAGTAAACTGGTTAGTAACTAA
ACTTCCAAACCAGTAAACTGGTTAGTAACTAAACTTCCAAACCAGTAAACTGGTTAGTAACTAAACTTCC
AAACCAGTAAACTGGTTAGTAACTAAACTTCCAAACCAGTAAACTGGTTAGTAACTAAACTTCCAAACCA
GTAAACTGGTTAGTAACTAAACTTCCAAACCAGTAAACTGGTTAGTAACTAAACTTCCAAACCAGTAAAC
TGGTTAGTAACTAAACTTCCAAACCAGTAAACTGGTTAGTAACTAAACTTCCAAACCAGTAAACTGGTTA
GTAACTAAACTTCCAAACCAGTAAACTGGTTAGTAACTAAACTTCCAAACCAGTAAACTGGTTAGTAACT
AAACTTCCAAACCAGTAAACTGGTTAGTAACTAAACTTCCAAACCAGTAAACTGGTTAGTAACTAAACTT
CCAAACCAGTAAACTGGTTAGTAACTAAACTTCCAAACCAGTAAACTGGTTAGTAACTAAACTTCCAAAC
CAGTAAACTGGTTAGTAACTAAACTTCCAAACCAGTAAACTGGTTAGTAACTAAACTTCCAAACCAGTAA
ACTGGTTAGTAACTAAACTTCCAAACCAGTAAACTGGTTAGTAACTAAACTTCCAAACCAGTAAACTGGT
TAGTAACTAAACTTCCAAACCAGTAAACTGGTTAGTAACTAAACTTCCAAACCAGTAAACTGGTTAGTAA
CTAAACTTCCAAACCAGTAAACTGGTTAGTAACTAAACTTCCAAACCAGTAAACTGGTTAGTAACTAAAC
TTCCAAACCAGTAAACTGGTTAGTAACTAAACTTCCAAACCAGTAAACTGGTTAGTAACTAAACTTCCAA
ACCAGTAAACTGGTTAGTAACTAAACTTCCAAACCAGTAAACTGGTTAGTAACTAAACTTCCAAACCAGT
AAACTGGTTAGTAACTAAACTTCCAAACCAGTAAACTGGTTAGTAACTAAACTTCCAAACCAGTAAACTG
GTTAGTAACTAAACTTCCAAACCAGTAAACTGGTTAGTAACTAAACTTCCAAACCAGTAAACTGGTTAGT
AACTAAACTTCCAAACCAGTAAACTGGTTAGTAACTAAACTTCCAAACCAGTAAACTGGTTAGTAACTAA
ACTTCCAAACCAGTAAACTGGTTAGTAACTAAACTTCCAAACCAGTAAACTGGTTAGTAACTAAACTTCC
AAACCAGTAAACTGGTTAGTAACTAAACTTCCAAACCAGTAAACTGGTTAGTAACTAAACTTCCAAACCA
GTAAACTGGTTAGTAACTAAACTTCCAAACCAGTAAACTGGTTAGTAACTAAACTTCCAAACCAGTAAAC
TGGTTAGTAACTAAACTTCCAAACCAGTAAACTGGTTAGTAACTAAACTTCCAAACCAGTAAACTGGTTA
GTAACTAAACTTCCAAACCAGTAAACTGGTTAG”

In one repeat cluster (position 349,933-350,462), the same imperfect palindrome repeat combines with a slightly different variant of the interspersed repeat, TAACTACTT, to constitute a composite repeat present in 14 tandemly arrayed copies.

“CAAACCAGTAAACTGGTTAGTAACAACTTCCAAACCAGTAAACTGGTTAGTAACAACTTCCAAACCAGT
AAACTGGTTAGTAACAACTTCCAAACCAGTAAACTGGTTAGTAACAACTTCCAAACCAGTAAACTGGTTA
GTAACTACTTCCAAACCAGTAAACTGGTTAGTAACAACTTCCAAACCAGTAAACTGGTTAGTAACAACTT
CCAAACCAGTAAACTGGTTAGTAACTACTTCCAAACCAGTAAACTGGTTAGTAACAACTTCCAAACCAGT
AAACTGGTTAGTAACAACTTCCAAACCAGTAAACTGGTTAGTAACAACTTCCAAACCAGTAAACTGGTTA
GTAACAACTTCCAAACCAGTAAACTGGTTAGTAACAACTTCCAAACCAGTAAACTGGTTAGTAACTACTT
CCAAACCAGTAAACTGGTTAGTAACAACTTCCAAACCAGTAAACTGGTTAGTAACAACTTCCAAACCAGT
AAACTGGTTAGTAACAACTTCCAAACCAGTAAACTGGTTAG”

Finally, the imperfect palindrome repeat “CAAACCAGT|AA|ACTGGTTAG” is also part of a 146 bp-long repeat that has 27 copies at different locations in the plastome. The sequence of this repeat unit is as follows:

“CTCGCTTGAACATATGTTCAAGCGAGCTAACTTTGCCTTGGCAAAGTTTGGTTATAAAGATAAGCCAGC
ATAGCTGACTTCCAAACCAGTAAACTGGTTAGTAACTAAACTTCCATATAATAGCTAAAGCTATTATAAG
TAACTGC”

**Note S4** Further details on the repeat insertions in the *L. pallida* plastid coding sequences

The open question existed of why the repeat units based on the imperfect palindrome (IP) motif AAGCCAGC|NNN|GCTGACTT and its variants are unevenly distributed in the direct and reverse complement orientation in exons. Our analysis for one particular version of the repeat unit, the “variant 4” repeat (summarised in Fig. 4d), shows that of the six possible reading frames (RFs) in which it may appear as part of the coding sequence, two (+1 and +2) contain a stop codon that would interrupt the coding sequence. Two RFs (-1 and -2) are potentially translatable, but do not occur in coding sequences of the *L. pallida* plastome. RF -1 would encode the two least frequent amino acids occurring in the *L. pallida* plastome-encoded proteins (C and W; see Table S6), one of them being encoded by the fourth rarest sense codon in the plastome (TGC; see Table S5). RF -2 also includes at least one infrequent codon (defined here as the sum of the least used codons together representing less than 4% of codons in the plastome, i.e. the 15 rarest codons from 61 sense codons; see Table S6). All the insertions found in the coding sequences correspond to RF +0 or -0, with the former much more abundant. None of them include codons of the rarest categories (2% or 4%), suggesting that they may be more efficiently translated than the insertions corresponding to RFs -1 and -2, and are thus better tolerated in the coding sequences. The dominance of RF +0 over RF -0 can also potentially be accounted for by the differences in the efficiency of translation, because RF +0 contains a lower number of codons categorised as “rare 10%” and “rare 20%” (see Fig. 4d; codons in these categories are listed in Table S6). Overall, it thus exhibits more optimal codon usage than RF -0. The effect of infrequent amino acids (W and C) and rare sense codons (below the 4% threshold) could also explain why the most abundant repeat in the plastome (CAAACCAGT|NN|ACTGGTTAG) neither broke into nor proliferated in the exonic regions (see Table S7).

**Note S5** Differential diagnosis of *Leontynka* spp. with regard to previously described colourless chlamydomonadalean taxa

Here we discuss all non-photosynthetic genera listed in Ettl [5] and based on the details provided by this most comprehensive monograph on green algal flagellates available. We preserve Ettl's classification of the genera to families, noting that the classification is outdated and does not reflect the actual relationships of the organisms.

<u>Dunaliellaceae:</u>
***Polytomella***. Fundamentally differs from *Leontynka* by possessing four flagella and lacking a cell wall. Moreover, starch is often stored only in the peripheral part of the lower two thirds of the cell in the form of small grains or slices. In contrast, starch grains in *Leontynka* are usually huge and occupying most of the cell's volume.
***Hyaliella***. The genus lacks papilla and an eyespot, while *Leontynka* has both.
***Hyalocardium***. Unlike *Leontynka*, cells of this genus are broadly heart-shaped, slightly flattened, clearly indented or hollowed-out in front, pointed or rounded at the rear, with the shape malleable.

<u>Asteromonadaceae:</u>
***Aulacomonas*** (=*Diphylleia* [14]). Morphologically distinct from *Leontynka* by its slightly quadrangular cells possessing a longitudinal furrow, giving the posterior end a bilobed appearance.
***Collodictyon.*** At first sight it differs from *Leontynka* by possessing four flagella. Moreover, the lateral cell margins are somewhat angular, with a broad, truncated but rounded apex, narrowing to the posterior, which may bear 1-3 lobes, or simply be broadly rounded. The cell posterior is often pseudopodial.
In fact, both *Diphylleia* and *Collodictyon* do not belong to the green algae, and constitute a clade belonging to the newly established eukaryotic supergroup CRuMs [15].

<u>Chlamydomonadaceae:</u>
***Polytoma***. *Leontynka* is generally similar to *Polytoma*, but does not match any of the previously described species. *Leontynka pallida* most resembles *P. papillatum* Pascher, described as having broadly ovoid (12.0-20.0 µm long and 9.0-13.0 µm wide) cells with a truncated papilla. However, *P. papillatum* does not have an eyespot. Meanwhile, *L. elongata* is generally similar to *P. eupapillatum* Ettl, a species that also has ellipsoidal to almost cylindrical cells, a keel-shaped papilla, and an ellipsoidal eyespot located in the lateral position of the second third of the anterior cell end. However, the cells of *P. eupapillatum* are slightly smaller (10.0-14.0 µm long and 5.0-6.0 µm wide) than those of *L. elongata*.
***Tussetia***. In contrast to *Leontynka*, this genus has a strongly tapered apex and does not accumulate starch.
***Hyalobrachion*****.** Significantly differs from *Leontynka* by having a spindle-shaped body with four short and thick rounded or pointed arms arising from the anterior half of the cell and oriented to the longer axis of the cell.

***Furcilia***. The genus shows a peculiar morphology with cells being strongly flattened, and broadly outlined from the base so that they consist of two large, lateral, backwardly curved, often somewhat unequally long and varyingly thick horns, which are held together in the middle by an almost spherical central part. This is very different from *Leontynka*.
***Tetrablepharis*.** Fundamentally differs from *Leontynka* by possessing four flagella.

Heamatococcaceae
***Hyalogonium*.** Distinguishable from *Leontynka* by having long spindle-shaped cells that are sharply pointed at both ends, but that are narrower at the rear than at the front.

Phacotaceae
***Chlamydoblepharis*.** In contrast to *Leontynka*, the genus contains a case, consisting of a coarse shell with a single apical flagellum opening, sometimes also with a thickened collar around the opening. Cells are decorated with fine pores or relatively large holes. None of these features were observed in *Leontynka*.

### Supplementary methods

**Methods S1** Isolation and cultivation of strains
The strain AMAZONIE was obtained from a freshwater hypoxic sediment sample collected near the “Explorer's Inn” lodge (12°50' S; 69°17' W), Tambopata province, Peru, in 2007. The strain MBURUCU was obtained from a freshwater hypoxic sediment sample collected near Mburucuyá, Corrientes province, Argentina, 28°02' S; 58°00' W, in 2013. The original samples were inoculated into 15 ml tubes containing 9 ml of Sonneborn’s *Paramecium* medium (ATCC medium 802 [16]); this medium was then used for routine cultivation. The cultures were kept at room temperature and subcultured once a week. The culture of AMAZONIE originally also contained *Paratrimastix* sp. (Metamonada: Preaxostyla), but became monoeukaryotic approximately a year after being established. The culture MBURUCU was never monoeukaryotic and contained *Trepomonas* sp. (Metamonada: Fornicata) besides *Leontynka*. Both cultures contained numerous unidentified bacteria.

**Methods S2** Light and transmission electron microscopy
Morphological investigations of the two colourless flagellates were conducted using an Olympus BX43 light microscope equipped with an Olympus DP27 digital camera (Tokyo, Japan). The Olympus micro imaging software cellSens v1.15 (Tokyo, Japan) was used for morphometric measurement of the algae. Vegetative cells were observed and documented using two-week-old cultures. One hundred cells of each organism were measured for size comparisons. For transmission electron microscopy (TEM), 900 µl of the cell suspensions of both AMAZONIE and MBUCURU were mixed with 100 µl of 25% (v/v) glutaraldehyde. Cells were aggregated into a pellet by centrifugation at 1000 g for 5 min at 12 °C. The supernatant was discarded and the remaining cell pellets were mixed with a fixative containing 2.5 % glutaraldehyde in a 0.1 M PIPES buffer. The specimens were then fixed in 1% (w/v) osmium tetroxide in a 0.1 M PIPES buffer on ice for one hour followed by dehydration through an ethanol series (30, 50, 70, 80, 90, 95, 100%) and substitution with acetone. The specimens were embedded in an Araldite Poly/Bed resin (Polyscience, Europe GmbH), which was polymerized at 70 °C for 48 hours. Ultrathin sections were cut on a Reichert-Jung Ultracut E ultramicrotome with a diamond knife, stained with uranyl acetate and lead citrate [17], and observed using a JEOL 1011 transmission electron microscope.

**Methods S3** Amplification and sequencing of 18S and ITS rDNA regions
Genomic DNA from both strains was extracted using the DNeasy Blood & Tissue Kit (Qiagen, Hilden, Germany) according to the manufacturer’s instructions. A genomic region corresponding to an almost complete 18S rRNA gene of the strain AMAZONIE was amplified by PCR with the MedlinA (5’-CTGGTTGATCCTGCCAG-3’) and MedlinB (5’-TGATCCTTCTGCAGGTTCACCTAC-3’) primers [18] using an annealing temperature of 55 °C. The PCR product was directly sequenced using the amplification primers, and a set of standard internal primers (577F, 577R, 1125F, 1055R, MedlinA [19]). The 18S rRNA gene from the strain MBURUCU was amplified using the 18S-F (5'-AACCTGGTTGATCCTGCCAGT-3' [19]) and vivi1650R (5'-TCACCAGCACACCCAAT-3' [20]) primers with an annealing temperature of 52 °C. These primers as well as 416-37R (5'-ATTTGCGCGCCTGCTGCCTTCC-3') and 895-916F (5'-GTCAGAGGTGAAATTCTTGGAT-3' [19]) were subsequently used for sequencing. The ITS2 region from both strains was amplified and

sequenced using the 1500bf (5'-GATGCATTCAACGAGCCTA-3' [21]) and ITSB (5'-CTTTTCCTCCGCTTATTGATATG-3' [22]) primers with an annealing temperature of 52 °C. PCR products were purified using the Zymoclean™ GEL DNA Recovery Kit (ZymoResearch, Irvine, USA) or the GenElute™ PCR Clean-Up Kit (Sigma, St. Louis, USA), and were directly sequenced with the standard Sanger dideoxy method.

**Methods S4** DNA and RNA isolation
To minimise the amount of bacteria in the cultures, a two-step filtration procedure described by Karnkowska *et al.* [23] was employed. First, the whole culture was resuspended and filtered through filter paper by gravity flow. Subsequently, the filtrate containing AMAZONIE cells was filtered through a 3 µm-pore polycarbonate filter (Whatman International Ltd., Maidstone, UK). The suspension of AMAZONIE cells remained in the medium above the filter, and bacteria were washed away by continuous addition of fresh medium (approximately two volumes of the original culture). To harvest the cells, the cell suspension from the second filtration step was transferred to a clean vessel and centrifuged at 1500 g for 10 min. Pelleted cells were then used for DNA or RNA extraction. Genomic DNA was extracted from 6 L of the culture using a modified CTAB/Phenol protocol recommended for *Chlamydomonas* (https://www.pacb.com/wp-content/uploads/2015/09/DNA-extraction-chlamy-CTAB-JGI.pdf); total RNA was isolated using the TRIzol Reagent (Life Technologies) with a standard extraction protocol followed by the Turbo DNA-free kit procedure (Invitrogen) to remove residual DNA contamination.
